# DNA-damaging bacteriocins from human *Escherichia coli* intestinal isolates trigger prophage induction and promote lysogeny

**DOI:** 10.64898/2026.05.26.727859

**Authors:** Caroline Henrot, Laurent Debarbieux, Marie-Agnès Petit

## Abstract

Lysogens - bacteria harbouring prophages - are prevalent in the human gut microbiota. Nevertheless, factors triggering induction or repression of prophages remain poorly characterized. Here, we studied the involvement of bacteriocins - antibacterials involved in bacteria-bacteria competition – in prophage induction. We screened a collection of 1,768 fecal *Escherichia coli* isolates for bacteriocin-producing strains and selected 30 to test their capacity to induce a λ-related coliphage. In these, we identified 74 bacteriocin genes and demonstrate that only those coding a DNA-damaging bacteriocin trigger prophage induction. From one strain co-producing an E-type endonuclease colicin and the Mcc1229 microcin, we demonstrate that these colicins induce a broad panel of temperate phage genera. Assessing bacterial competition by pairwise cocultures between an E-type endonuclease-producing strain and a λ-lysogen revealed enhanced prophage induction and increased emergence of new lysogens among the bacteriocin producers. Remarkably, while the λ-lysogen was outcompeted by the E-type endonuclease colicin producer within 6 hours *in vitro*, both populations were maintained at comparable levels over 10 days in dixenic mice. This work reveals a dual role for DNA-damaging bacteriocins that kill competitors by prophage induction and propagate lysogeny.

## Introduction

The mammalian gut microbiota comprises diverse microbes among which bacteria are the most abundant and extensively studied, in contrast to bacteriophages (phages), viruses infecting bacteria, that remain understudied. Most phages display either a virulent (strictly lytic) or a temperate lifestyle. Importantly, in the gut, about 70 to 80% of the bacteria are lysogens that carry temperate phage genomes (prophages) as episome or integrated into their chromosome. Free phage particles are also abundant in the human gut, with estimated concentrations of 10^9^-10^10^ viral-like particles (VLPs) compared to approximatively 10^11^ bacteria per gram of stool [1–3]. Therefore, phages are suspected to shape intestinal bacterial communities through interactions ranging from predation by virulent phages, lysis by prophage induction, lysogeny and horizontal gene transfers by transduction.

DNA-damaging agents, such as mitomycin C (MmC) or ciprofloxacin, are classical triggers of prophage induction *in vitro*, leading to the production of new phage particles and their release by host lysis. In the typical case of *Escherichia coli* strains carrying phage lambda, DNA-double strand breaks activate RecA, further leading to the destabilization of the lambda repressor, parallel to the activation of LexA that regulates the SOS response [4]. Beyond this well characterized RecA-dependent mechanism, other RecA-independent mechanisms have been described [5–7]. Nevertheless, for the well-known *E. coli* Mu and P2 prophages, triggering factors and associated mechanisms have not yet been identified.

In the gut, triggering factors can originate either from external uptakes, from the host metabolism or the microbiota itself [8]. Diet enriched in fructose enhanced acetic acid production, which activated *Lactobacillus reuteri* stress response, leading to enhanced prophage induction [9]. Colibactin increases *in vitro* the induction of prophages from a broad spectrum of intestinal bacteria [10]. Recently, human host-associated cellular products were also identified *in vitro* as prophage induction triggers [11]. Moreover, our previous study on prophage activity in the isobiotic mouse line OMM^12^ demonstrated that at least one prophage is repressed *in vitro* while it is well induced in the gut [12].

Bacteriocins, which are ribosomally synthesized antimicrobials with narrow spectrum and often encoded on plasmids, are also putative triggering factors, with four of them reported to induce Stx-prophages *in vitro* [13,14]. Two types of bacteriocins are distinguished for *Enterobacteriaceae*: colicins (40-80 kDa) and microcins (<10 kDa). Notably, it was reported that in the genome of 54 % of human fecal *E. coli* isolates at least one colicin or microcin gene cluster were identified, with higher frequencies in the B2 phylogroup [15]. While microcins are systematically secreted by bacteria, colicins are released upon host lysis. Both bacteriocins display various activities. Most of colicins are pore-forming molecules, others degrade nucleic acids - targeting DNA, mRNA or specific tRNAs - and less frequently, some inhibit peptidoglycan synthesis [16]. Microcins activities spans from disrupting membrane integrity, protein synthesis or DNA and RNA metabolism [17,18]. Regulation of colicin production is typically triggered by LexA upon DNA damages [19], and it was reported that the synthesis of some colicins and microcins is increased upon nutrient depletion and in stationary growth phase [20–22]. Moreover, many microcin clusters are under the regulation of Ferric Uptake Regulator (Fur) and thus increasingly produced in iron-limited conditions including in the gut [23–25].

To evaluate the prevalence of bacteriocins and their putative triggering activity on prophage induction, we leveraged a large collection of fecal *E. coli* isolates originating from a cohort of one year-old children (COPSAC) [26,27]. From this collection, we previously characterized the lysogens and showed that induced prophages are mostly related to phages lambda and P2 [28]. First, we screened 1,768 isolates for bacteriocin-producing strains that do not produce spontaneously phage particles and second, we tested 30 representative candidates for their ability to induce prophages of lysogens from the same collection. We found that supernatants of more than 20% of colicin-producers induce prophages. Moreover, when introduced in an *E. coli* MG1655 derivative, one of these colicins provided an advantage *in vitro* over an isogenic strain carrying a lambda prophage by boosting prophage induction and lysis while also promoting lysogenization. Interestingly, when these two strains were introduced in dixenic mice, the lambda lysogen was no longer outcompeted despite the production of colicin, suggesting that the intestinal environment modulates their competitive dynamics.

## Results

### Most of *Escherichia coli* isolates from the COPSAC collection spontaneously release antibacterials and/or phage particles

To evaluate the release of antibacterials by 1,768 *E. coli* COPSAC isolates, we performed spot-on-lawn assays using as indicator strains either the *E. coli* strain C or the strain MG1655 *hsdR* that is defective for the EcoKI restriction modification system [28] (**Fig 1**). This qualitative method, commonly used to detect bacteriocin-producing strains, translates into zones of inhibition (ZOI) around the spots [29]. Strikingly, 65% (1,159 out of 1,768) of the isolates produced ZOI on at least one indicator strain (**Table 1**). To determine to which extent prophage induction contributed to this pattern, filtered supernatants were streaked on agar plates and then covered by a bacterial lawn (strain C or MG1655 *hsdR*) (**Fig 1**). Spontaneous phage release was observed for 74% (859 out of 1,159) of ZOI-positive strains and 39% (241 out of 609) of ZOI-negative isolates, showing that prophage induction was not strictly associated with ZOI (**Table 1**). Overall, 17% (300 of 1,768) of isolates formed ZOI without spontaneously releasing phages. We conclude that the release of antibacterials is highly prevalent among *E. coli* fecal isolates (79% of the collection, i.e. all strains but the 368 for which no ZOI nor spontaneous phage release were observed suggestive of dynamic intraspecies competitions in the gut microbiota when several strains of *E. coli* co-occur [30].

**Figure 1.**
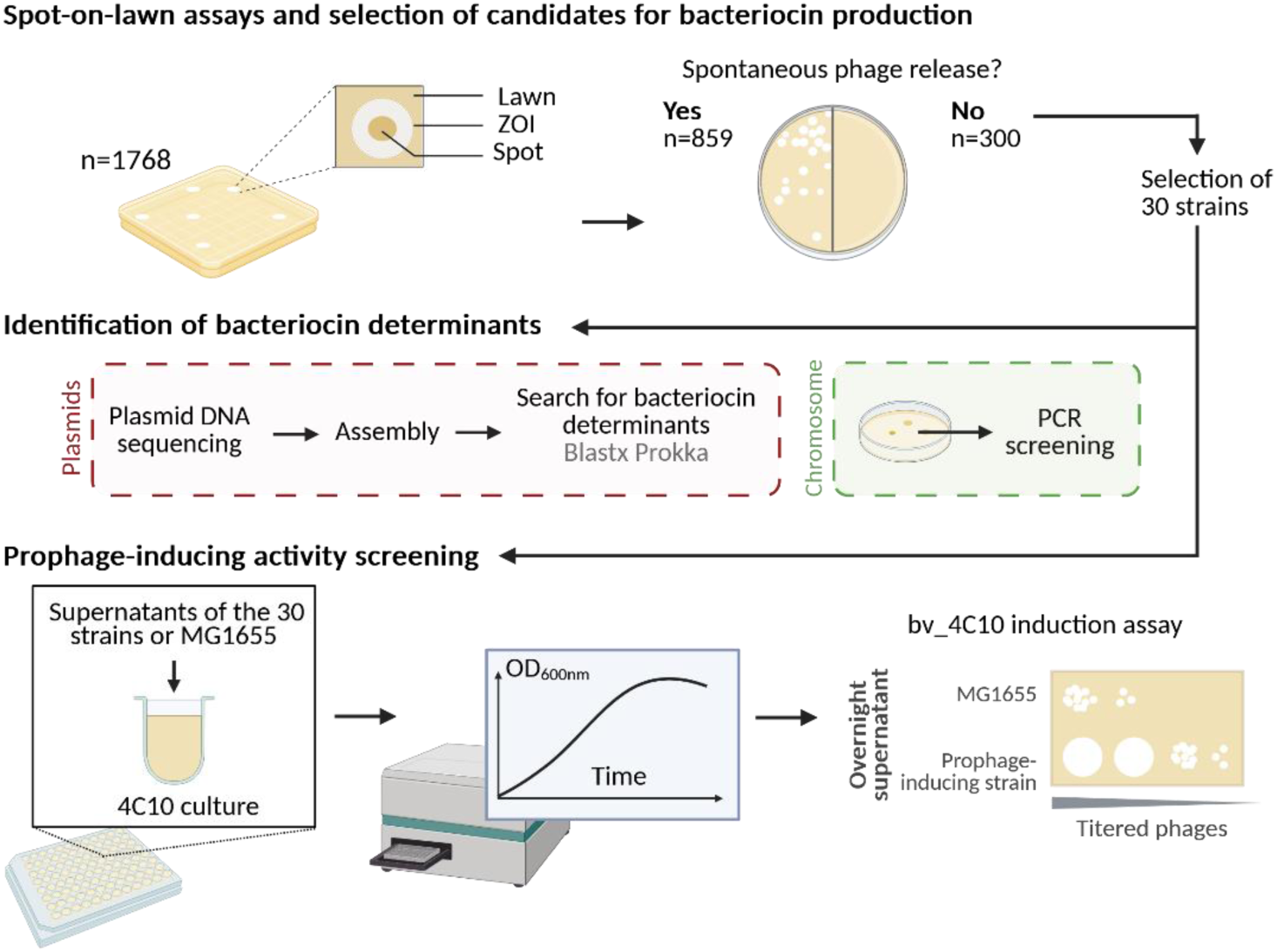
Scheme presenting the method to identify and analyse strains producing prophage-inducing bacteriocins. From 1,768 *E. coli* fecal isolates, 30 strains producing Zone of Inhibition (ZOI) without spontaneous phage release were randomly selected, and their plasmid and bacteriocin gene content were determined. In parallel, the ability of their supernatants to induce a lambda-related prophage (bv_4C10) from the lysogen 4C10 was evaluated. Created in BioRender. Henrot, C. (2026) https://BioRender.com/frd0zh0

**Table 1.**
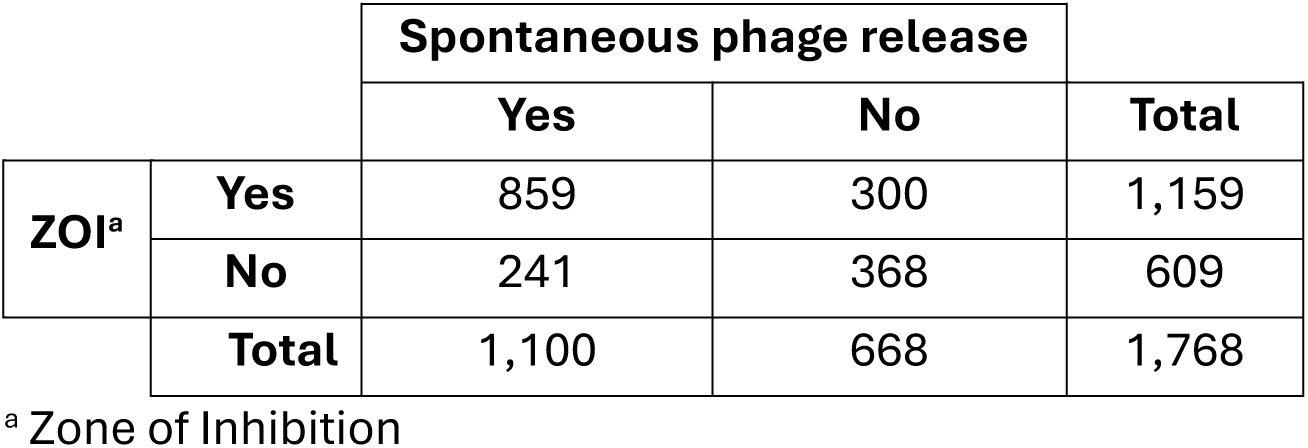
Number of ZOI-positive and phage-releasing COPSAC *E. coli* isolates.

### *E. coli* gut isolates are rich in bacteriocin encoding genes

Among the above 300 *E. coli* COPSAC isolates positive for ZOI and negative for spontaneous prophage induction, we randomly selected a set of 30 and assess their content in bacteriocin-encoding genes (**Fig 1**). These 30 isolates displayed a diversity of ZOI diameters and aspects (**Fig 2A**). Among them, 24 contain one to six plasmids that were deep-sequenced and assembled (**S1A Fig**). To identify bacteriocin-encoding genes, automatic Prokka annotations of all assembled contigs were complemented by blastx searches against an in-house database of *Enterobacteriaceae* bacteriocin proteins (see Methods, **S1** and **S2 Tables**). In addition, for each of the 30 isolates, we assessed by PCR with specific primers the presence of the two chromosomally encoded microcins, MccM and MccH47 [15]. Interestingly, for five of the six PCR positive strains that host plasmid(s), the presence of one or bothmicrocin genes was confirmed among the residual chromosomal contigs assembled from the sequenced plasmids. Overall, we identified a total of 74 bacteriocin-encoding genes belonging to 18 different types out of the 38 included in our database (**Fig 2B; S1B Fig; Supplementary information**). In total, only two isolates (4H1 and 15C2) do not carry bacteriocin genes, while the 28 others possess either one or several genes (**S1C Fig**), which illustrates the richness in bacteriocin determinants among these 30 isolates. We next tested the capacity of supernatants from these 30 isolates to induce prophages.

**Figure 2.**
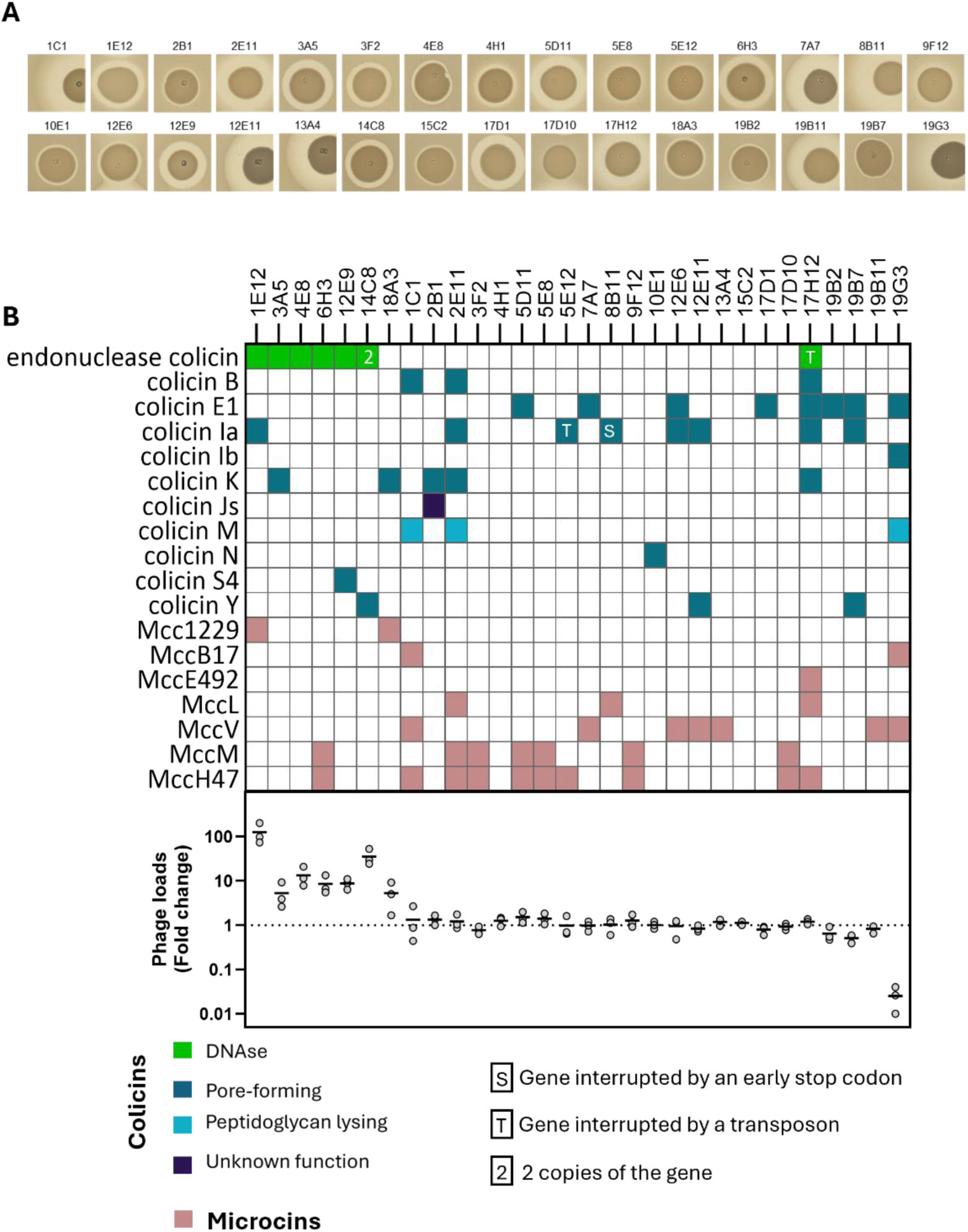
Prophage-inducing supernatants originate from *E. coli* isolates encoding an E-type endonuclease colicin or microcin Mcc1229. **A.** Representative pictures of zone of inhibition (ZOI) from spot-on-lawn assays, on lawns of strain C for the 30 *E. coli* isolates selected to test bacteriocin production and prophage-inducing capacity. **B.** The top plot lists bacteriocin genes identified in the 30 *E. coli* isolates, among which only endonuclease colicins and Mcc1229 are associated with prophage induction. The bottom plot reports the induction level of *Bievrevirus* bv_4C10 by the corresponding strain supernatants normalized to the concentration of phage spontaneously produced when strain 4C10 is treated with spent medium supernatant of strain MG1655 (∼5.5E6 PFU.mL^-1^). Each dot represents one experiment (n=3 independent experiments).

### E-type endonuclease colicins and Mcc1229 microcin are strongly associated with prophage induction

To detect prophage induction by bacteriocins, we used the *E. coli* COPSAC isolate 4C10 that hosts a lambda-related prophage assigned to the *Bievrevirus* genus (Genbank LR595861) whose master regulator is most similar to that of phage P22 [28]. Particles of the induced prophage bv_4C10 were titrated on *E. coli* strain C. First, we evaluated the putative toxicity of bacteriocins on strain 4C10. A liquid culture of 4C10 in early exponential phase was mixed with an equal volume of filtered supernatants from overnight liquid cultures of each of the 30 *E. coli* isolates and 4C10 growth was monitored during 180 min (**Fig 1**). Strikingly, 4C10 growth was largely unaffected except by the supernatant from isolate 19G3 that encodes the well-known colicin Ib [31] (**S2 Fig**).

Second, supernatants of the above 4C10 cultures collected at 180 min after exposure to either the 30 isolates supernatants or a spent medium from a culture of the *E. coli* strain MG1655 used as control, were titrated on strain C (**Fig 1**). A total of seven (23%) supernatants led to a phage titre increase from 5 to 120-fold (**Fig 2A**). Strikingly, six of these supernatants originated from the strains carrying a complete endonuclease colicin cluster (1E12, 3A5, 4E8, 6H3, 12E9 and 14C8). Another supernatant from the isolate 17H12 that encodes an endonuclease colicin cluster did not induce bv_4C10, likely because a transposon interrupts the endonuclease gene, which consolidates a strong association between endonuclease colicins and bv_4C10 induction (**S1D Fig**).

The seventh prophage-inducing supernatant originated from isolate 18A3, carrying one copy of colicin K and microcin Mcc1229. Since colicin K is also present in 3 strains (2B1, 2E11, 17H12) from which supernatants did not induce bv_4C10, we concluded that Mcc1229 was likely involved in prophage induction from strain 18A3. Moreover, we noted that the isolate 1E12, which carries genes for an E-type endonuclease colicin and the Mcc1229 on two different plasmids, led to the highest titre of bv_4C10, suggesting that this elevated induction results from a cumulative effect of these two bacteriocins.

### Both microcin Mcc1229 and E-type endonuclease colicin induce bv_4C10

We next examined the individual contribution of the E-type endonuclease colicin and microcin Mcc1229 from strain 1E12, encoded on plasmids p1_1E12 and p2_1E12, respectively (**S3 Fig**). Following the insertion of a kanamycin resistance cassette in each plasmid (resulting in plasmids pEndo and pMcc1229, respectively) and their subsequent transformation into strain MG1655 (see **Tables 2** and **3**), we confirmed that both bacteriocins were active as they formed ZOI on strain C, while control strains carrying plasmids in which bacteriocin genes were deleted (pΔEndo and pΔMcc1229) did not (**Fig 3A**).

**Figure 3.**
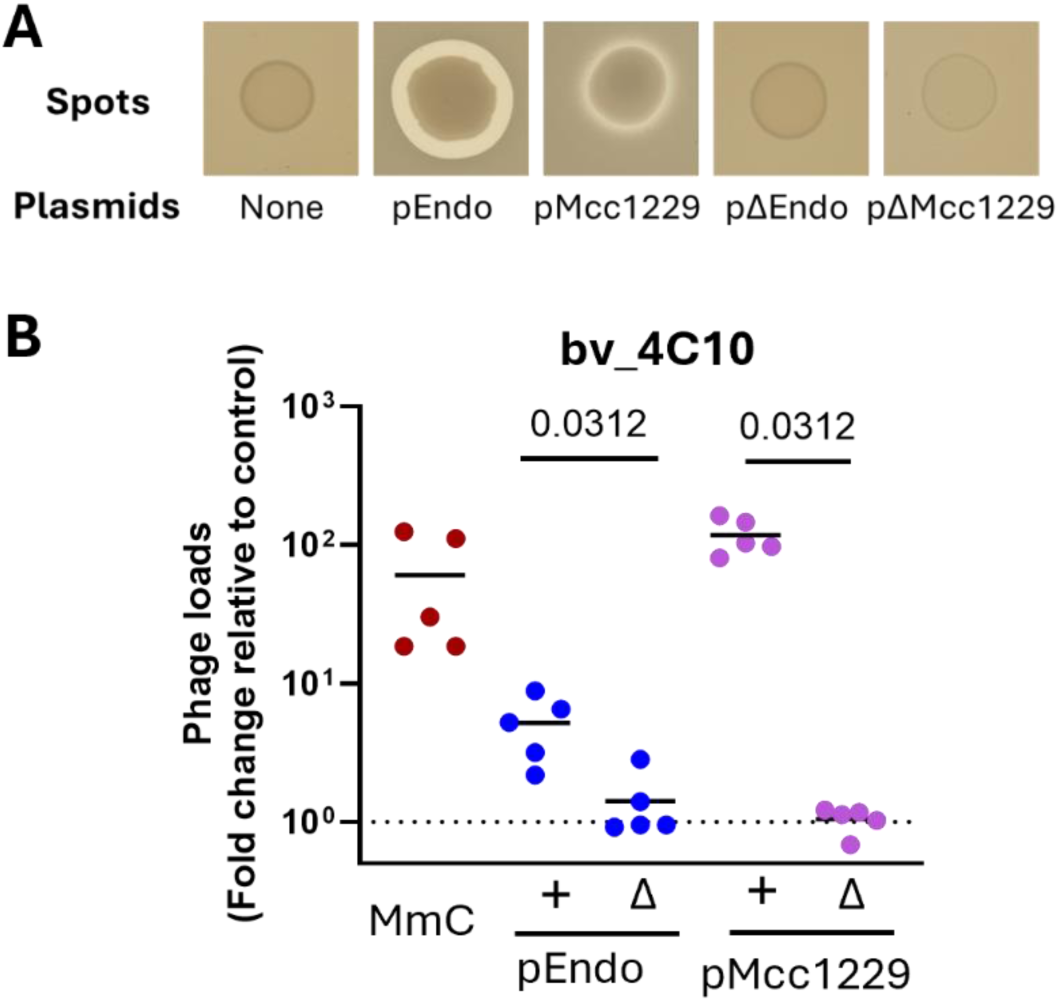
Microcin Mcc1229 and E-type endonuclease colicin independently induce the *Bievrevirus* bv_4C10. **A.** Spot-on-lawn assays on the C strain with spots from the MG1655 strain transformed with either pEndo or pMcc1229 and their respective inactive variants pΔEndo or pΔMcc1229. **B.** Supernatants of the above strains were incubated with strain 4C10 during 180 min and then plated on strain C to count PFUs of the induced *Bievrevirus* bv_4C10. Results are expressed relative to the control (MG1655 without plasmid) with n=5 independent experiments. One-tail Wilcoxon tests for paired samples revealed a significant difference between pEndo (or pMcc1229) and pΔEndo (or pΔMcc1229) conditions.

**Table 2.**
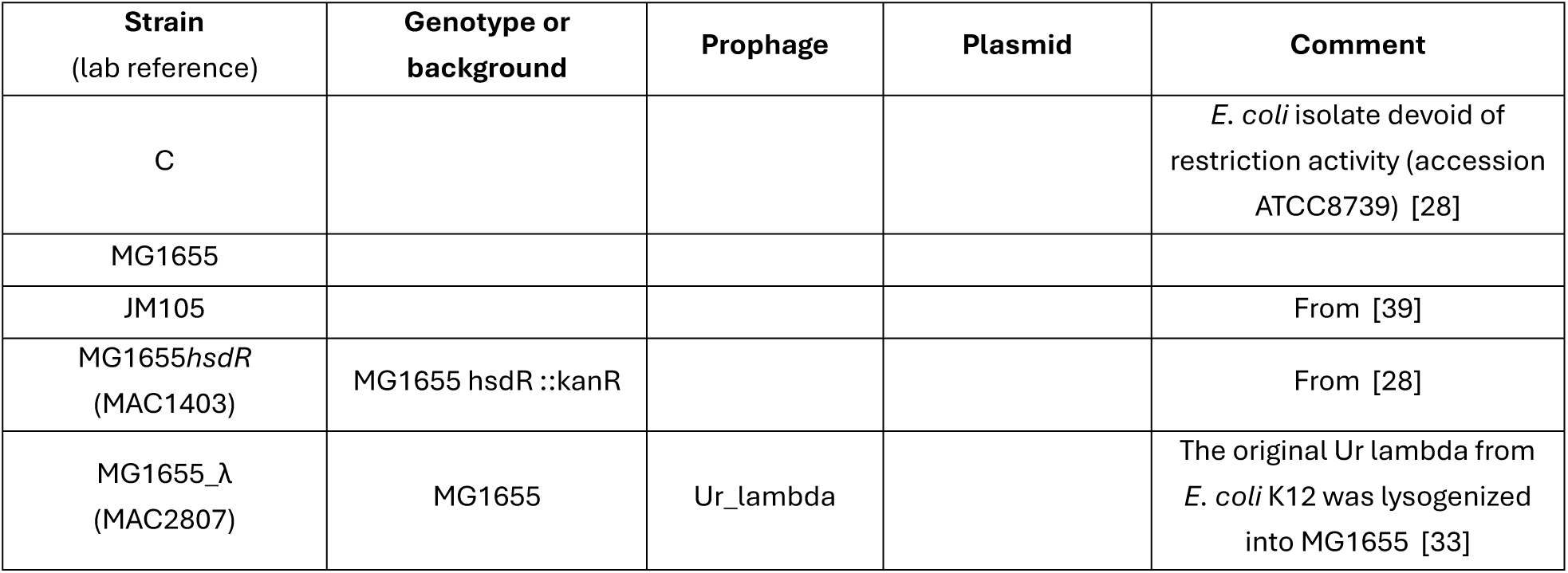
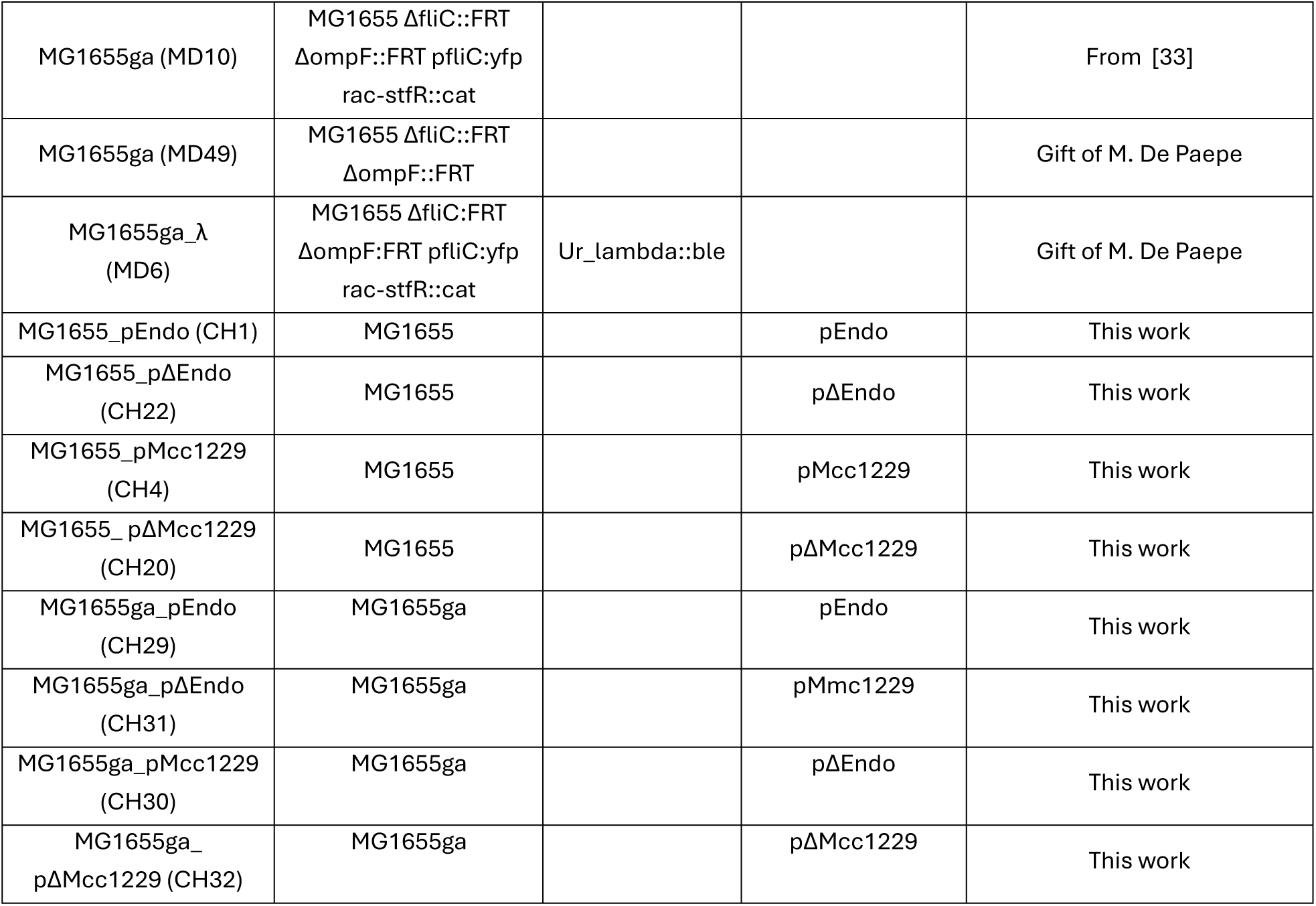
List of the strains used in this study.

| Strain<br>(lab reference) | Genotype or<br>background | Prophage | Plasmid | Comment |
| --- | --- | --- | --- | --- |
| C |  |  |  | *E. coli* isolate devoid of restriction activity (accession ATCC8739) [28] |
| MG1655 |  |  |  |  |
| JM105 |  |  |  | From [39] |
| MG1655*hsdR*<br>(MAC1403) | MG1655 *hsdR* ::kanR |  |  | From [28] |
| MG1655_λ<br>(MAC2807) | MG1655 | Ur_lambda |  | The original Ur lambda from *E. coli* K12 was lysogenized into MG1655 [33] |
| MG1655ga (MD10) | MG1655 ΔfliC::FRT<br>ΔompF::FRT pfliC::yfp<br>rac-stfR::cat | | | From [33] |
| MG1655ga (MD49) | MG1655 ΔfliC::FRT<br>ΔompF::FRT | | | Gift of M. De Paepe |
| MG1655ga_$\lambda$<br>(MD6) | MG1655 ΔfliC::FRT<br>ΔompF::FRT pfliC::yfp<br>rac-stfR::cat | Ur_lambda::ble | | Gift of M. De Paepe |
| MG1655_pEndo (CH1) | MG1655 |  | pEndo | This work |
| MG1655_pΔEndo<br>(CH22) | MG1655 | | pΔEndo | This work |
| MG1655_pMcc1229<br>(CH4) | MG1655 |  | pMcc1229 | This work |
| MG1655_pΔMcc1229<br>(CH20) | MG1655 | | pΔMcc1229 | This work |
| MG1655ga_pEndo<br>(CH29) | MG1655ga |  | pEndo | This work |
| MG1655ga_pΔEndo<br>(CH31) | MG1655ga | | pMcc1229 | This work |
| MG1655ga_pMcc1229<br>(CH30) | MG1655ga | | pΔEndo | This work |
| MG1655ga_<br>pΔMcc1229 (CH32) | MG1655ga | | pΔMcc1229 | This work |

**Table 3.** Plasmid constructions.

| Plasmids | Plasmid description | Insert |  |  |  | Vector |  |
| --- | --- | --- | --- | --- | --- | --- | --- |
|  |  | Gene amplified by PCR | DNA matrix | PCR oligo 1 | PCR oligo 2 | Original plasmid | Digested by |
| pEndo | kanR | kanR | pSMG261 | OPM60 | OFL368 | p1_1E12 | PmlI |
| pΔEndo | kanR ΔceaE (1975-3591) |  |  |  |  | pEndo | DraI |
| pMcc1229 | kanR | kanR | pACYC177 | OPM60 | OFL368 | p2_1E12 | BamHI |
| pΔMcc1229 | kanR ΔmctA (1851-2191) |  |  |  |  | pMcc1229 | BmtI + EcoRI |

Next, we quantified bv_4C10 induction by the supernatants of four isogenic MG1655 strains carrying each of the above plasmids, taking as a reference the induction by the supernatant of an MG1655 strain without plasmid. The supernatant of MG1655_pΔMcc1229 did not increase the level of bv_4C10 particles compared to the control. In sharp contrast, the supernatant of MG1655_pMcc1229 resulted in a 112-fold increase of bv_4C10 particles (p= 0.0312, One-tail Wilcoxon test for paired samples), reaching a higher level than MmC-triggered induction (**Fig 3B**). Interestingly, this elevated induction was associated with a growth inhibition of strain 4C10, while MmC induction was not (**S4A Fig**). The E-type endonuclease colicin also triggered the induction of bv_4C10 (4-fold, p= 0.0312, One-tail Wilcoxon test for paired samples), albeit to a lesser extent than Mcc1229 (**Fig 3B**). Therefore, the two bacteriocins from isolate 1E12 can independently trigger the induction of bv_4C10.

### The microcin Mc1229 and the E-type endonuclease colicin induce taxonomically distinct prophages but not *Peduoviruses*

Moving forward, we evaluated the capacity of these two bacteriocins to induce prophages distinct from bv_4C10. First, we tested the most studied *E. coli* prophage, Ur_lambda, belonging to the *lambdavirus* genus, which master regulator CI is distinct from the P22-like repressor of phage bv_4C10. Ur_lambda was lysogenized into strain MG1655 (strain MG1655_λ, **Table 2**). Supernatants from strains MG1655_pEndo and MG1655_pMcc1229 resulted in a drastic growth inhibition of strain MG1655_λ (**S4B Fig**) and an increased phage particle release, compared to their controls (p= 0.0312, One-tail Wilcoxon test for paired samples, **Fig 4A**). Second, we selected several lysogens among the 1,768 intestinal *E. coli* isolates hosting other prophages distantly related to lambda (*Bievrevirus* in 4A7, *Limmatquaivirus* in 4F5, the latter encoding a *S. enterica* LexA type of master regulator distinct from that of bv_4C10 or lambda), as well as isolates hosting *Peduovirus* genus prophages (2H4, 4B2, 4C9, 4E6) [28] (**Table 2**). These lysogens were exposed to the supernatants from three strains - 1E12 or MG1655_pEndo or MG1655_pMcc1229 - and the identity of induced prophages was confirmed by PCR from plaques.

**Figure 4.**
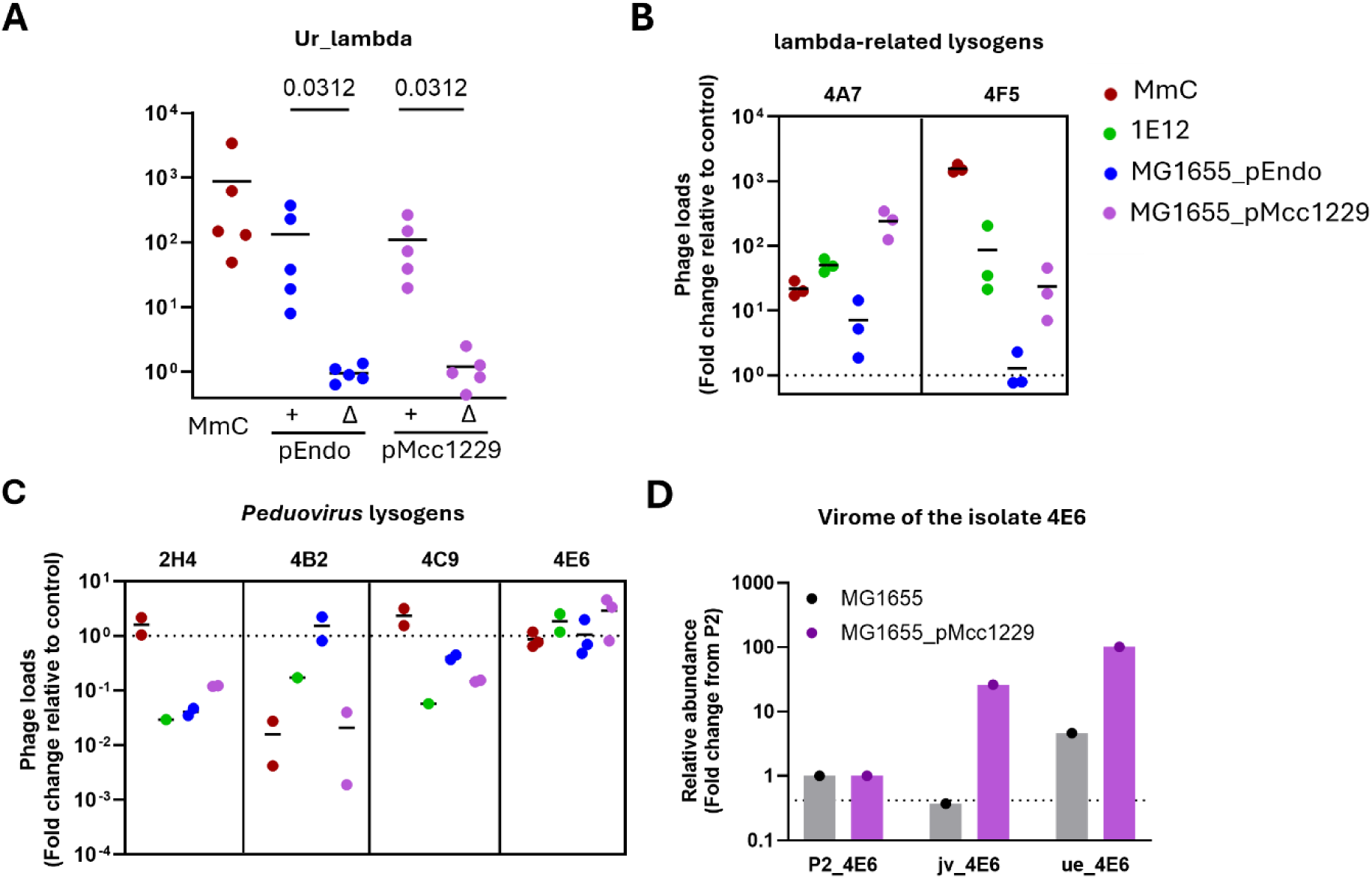
E-type endonuclease and Mcc1229 induce prophages from different genera. **A.** Induction of Ur_lambda prophage by the supernatants of strains MG1655 carrying the indicated plasmids were plotted relative to the MG1655 plasmid-less control (n= 5 independent experiments). One-tail Wilcoxon tests for paired samples were significant comparing pEndo (or pMcc1229) and pΔEndo (or pΔMcc1229) conditions. **B.** Induction of lambda-related prophages (from *Limatquaivirus* and *Jouyvirus* genera) is reported as in panel A. **C.** Induction of *Peduovirus* prophages is reported as in panel A. Reduction of phage loads is linked to a lower bacterial density upon treatment. **D.** Abundance of reads mapping to P2, jv_4E6 and ue_4E6 phage genomes, using the viromes of strain 4E6 exposed to supernatants of indicated strains. Dashed line represents the relative abundance of bacterial contigs for the lysogen exposed to MG1655 supernatant.

Exposure of lysogens 4A7 and 4F5 to MG1655_pMcc1229 or 1E12 supernatants resulted in a growth defect after 80 min, suggesting extensive prophage induction (**S4C Fig**), which was corroborated by a 2-log increase of phage titres for *Bievrevirus* 4A7 and *Limatquaivirus* 4F5 (**Fig 4B**). Increased PFUs were also observed following exposure to supernatant of MG1655_pEndo for 4A7 but not for 4F5. In contrast, when lysogens hosting *Peduovirus* prophages were exposed to each of the three supernatants, we did not observe an increase of P2 phage titres compared to the control condition (**Fig 4C**). Nevertheless, growth of these lysogens was affected after 60 minutes for strains 4E6, 4C9 and 2H4 exposed to Mcc1229-containing supernatants, suggesting that some other prophages could be induced but remained undetected by spot assay on the lawn of strain C (**S4C Fig**). To identify these putative prophages, DNA viromes from these strains exposed to either MG1655 or Mcc1229 supernatants were deep-sequenced. In strain 4E6 supernatants, two phages were identified besides the expected *Peduovirus.* One, named ue_4E6, belongs to the *Uetakevirus phiV10* species [32]. The other, named jv_4E6, is a lambda-like from the *Jouyvirus* genus (encoding a master regulator most similar to the one of phage 933W). To estimate their induction levels, reads were mapped back on the phage genomes and reads coverage of each phage was normalized to reads mapping on the *Peduovirus,* which concentration, based on plaque assays, remained globally stable across conditions (**Fig 4C**). Upon microcin treatment, coverage ratios were 21- and 70-fold higher for ue_4E6 and jv_4E6, respectively, compared to the P2 control, showing that Mcc1229 induces both prophages (**Fig 4D**). Using the same approach, a *Uetakevirus* phage from 4C9 (ue_4C9) and a lambda_like phage from 2H4 (Josas_2H4, with an *E. coli* LexA type of master regulator) were also found induced (47 and 89-fold, respectively) upon exposure to Mcc1229 (**S4D Fig**).

Therefore, we conclude that the two bacteriocins of strain 1E12 induce prophages from the *Bievrevirus* and *Lambdavirus* genera, which encode distinct repressors, and that at least the microcin Mcc1229 also induces prophages from the *Limatquaivirus*, *Uetakevirus* and *Jouyvirus* genera, as well as the distinct Josas_2H4 prophage. Interestingly, the induction of *Peduovirus* prophages were not enhanced by any of these two bacteriocins.

### *E. coli* producing the E-type endonuclease colicin triggers prophage induction and outcompetes an isogenic lysogen *in vitro*

Next, we sought to evaluate whether prophage induction by bacteriocin could affect bacteria-bacteria competition *in vitro* and in the gut of mice. For this, we used the strain MG1655ga, which carries *fliC* and *ompF* mutations that were previously reported to prevent the fixation of *ompB* mutations in strain MG1655 during the colonization of germ-free mice, which in turn lower the levels of phage lambda receptor LamB [33].

We then used a lysogen susceptible to bacteriocins in the same genetic background with a chloramphenicol resistance marker and carrying the Ur_lambda prophage modified to encode a phleomycin resistance marker (MG1655ga_λ). This phleomycin marker allows for the detection of putative new lysogens [33]. Indeed, we anticipated that mixing the strains MG1655ga_λ with MG1655ga_pEndo would release Ur_lambda particles that could either replicate (free phage titrated on strain C) or lysogenize the susceptible MG1655ga_pEndo strain (**Fig 5A**).

**Figure 5.**
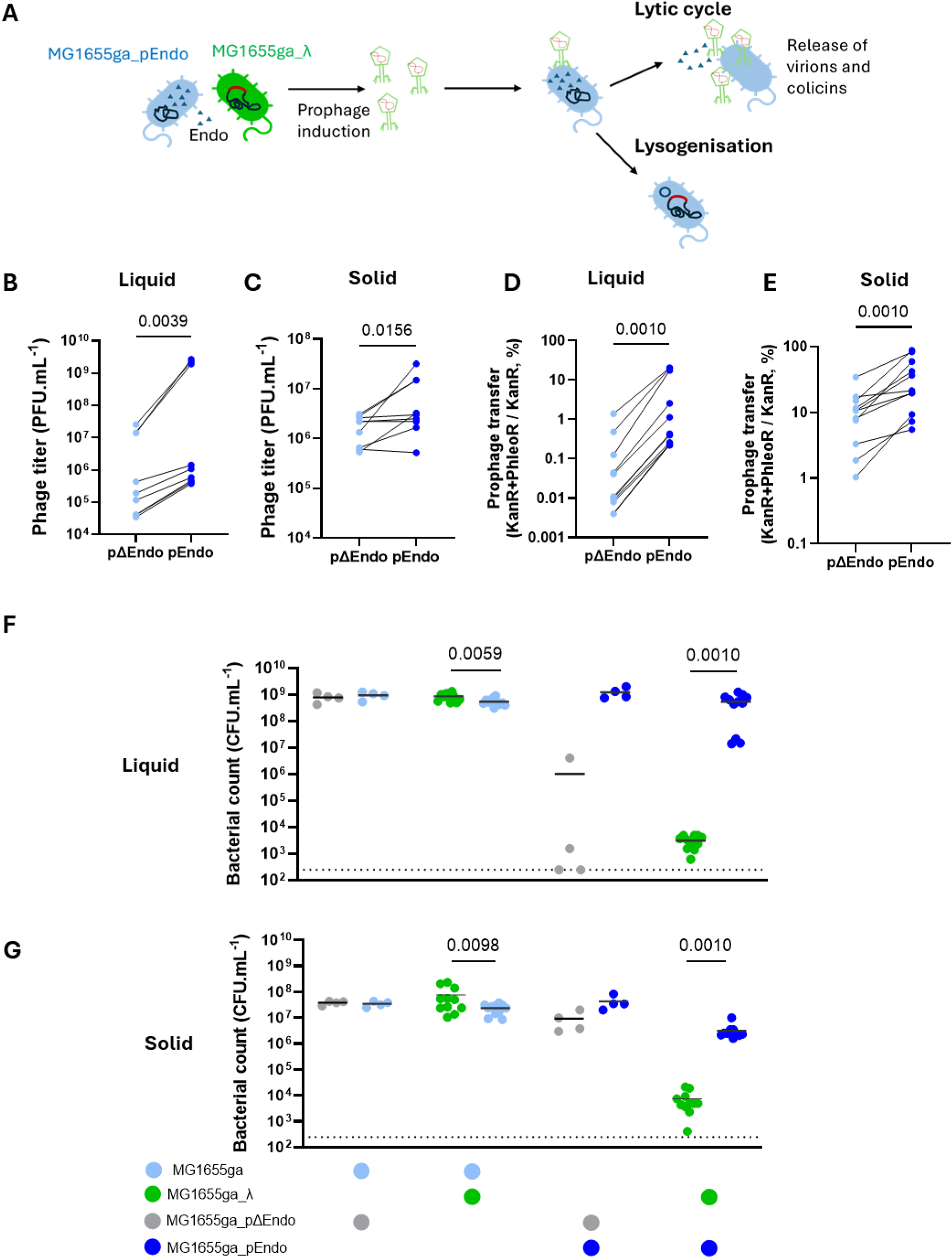
A strain carrying the E-type endonuclease colicin is lysogenised by the prophage it induces from an isogenic strain. **A.** Schematic illustration of expected events during the competition between two isogenic strains, one carrying the E-type endonuclease colicin and the other the lambda prophage. **B-G.** Equal volumes of strains MG1655ga_λ (CmR, PhleoR) or MG1655ga (CmR) were mixed with strains MG1655ga_pEndo (KanR) or MG1655ga_pΔEndo (KanR) and incubated in well mixed (“liquid”) or on agar (“solid”) conditions during 6 hours before assessing free phages particles (**B,C**) on C strain, prophage transfer by lysogenization of Ur_lambda::ble (**D,E**) on selective medium and bacterial densities in each mix (**F,G**) including the proportion of newly lysogenized bacteria, with appropriate markers. Individual values (n=4 to 11 independent experiments) are represented with the mean (horizontal bars; p values were obtained from Wilcoxon test for paired samples).

To perform competition experiments *in vitro,* strains MG1655ga_λ and MG1655ga_pEndo (or MG1655ga_pΔEndo as a control) were grown to exponential phase, mixed in equal ratio, and propagated for 6 h in tubes (well-mixed condition hereafter designated as “liquid”) and agar plates (designated as “solid”), to compare conditions with contrasted bacteriocin diffusion and bacterial contacts (see Methods). After 6 h of co-incubation in the liquid condition, free phage loads were significantly increased when the strain MG1655ga_λ was mixed with MG1655ga_pEndo compared to MG1655ga_pΔEndo (p=0.0039, Wilcoxon test for paired samples). Despite higher variability between replicates, significant increase was also observed in the solid condition at 6 h (p= 0.0156, **Figs 5B-C**). The increase of new lysogens (MG1655ga_pEndo or MG1655ga_pΔEndo lysogenized by λ) at 6 h was significant in both conditions (p= 0.0010, **Figs 5D-E**).

We next evaluated how prophage induction affected the proportion of the two strains in competition. MG1655ga_pΔEndo co-cultured with MG1655ga resulted in similar bacterial counts, whereas MG1655ga_pΔEndo with MG1655ga_λ led to a significant 1.5-fold advantage for the lysogen in liquid (p=0.0059) and a 4-fold advantage in solid (p=0.0098) conditions (Wilcoxon test for paired samples), thereby showing a moderate but significant fitness advantage likely due to spontaneous prophage induction (**Figs 5F-G**). By contrast, counts of MG1655ga_λ were significantly reduced by 5- and 3-log when co-cultured with MG1655ga_pEndo in liquid (p=0.0010) and solid (p=0.0010) conditions, showing that the colicin-producing strain outcompeted the lysogen.

To determine to which extent lysogeny increased susceptibility to the colicin-producing strain (MG1655ga_pEndo), we compared MG1655ga_λ with its isogenic prophage-less strain MG1655ga. In the liquid condition, MG1655ga density was strongly reduced with two replicates in which this strain was undetectable, a behaviour similar to the lysogen (**Fig 5F**), whereas in solid, MG1655ga was barely affected by the colicin (**Fig 5G**).

Finally, similar experiments were conducted with pMcc1229. After 6 h of co-culture, Mcc1229 had a significant effect on prophage transfer, both in liquid and solid conditions, as observed for the endonuclease (**S5 Fig**). However, the free phage titres were significantly increased only in the solid condition. This differs from the endonuclease effect, where free phages were increased in both conditions, suggesting that the microcin is more effective when bacteria grow in close contact. In line with these observations, the microcin-producing strain did not outcompete the lysogen in liquid in contrast to the solid condition. However, this effect was independent of the presence or absence of a prophage in the competing strain (**S5 Fig**).

Altogether, competition experiments revealed that the strain expressing the E-type endonuclease colicin displayed broader effects than the strain expressing the microcin. Indeed, the endonuclease triggered Ur_lambda induction in liquid condition, boosted its lysogenization and, on solid condition, outcompeted the lysogen, all to a greater degree than the isogenic non-lysogen. These results encouraged us to assess its activity in the gut of mice.

### The E-type endonuclease colicin does not increase prophage induction in the mice gut

We then used germ-free mice to assess the potential of the E-type endonuclease colicin to increase prophage dynamics in the gut. Equal amounts of strains MG1655ga_λ and either MG1655ga_pEndo or MG1655ga_pΔEndo were given to mice in two independent experiments, during which animals exhibited no signs of distress. Free phage titres, lysogenization frequencies and bacterial loads were estimated daily in fecal samples and in gut sections (luminal and mucosal contents of both ileum and colon) at the end point (10 days post-gavage) (**Fig 6A**).

**Figure 6.**
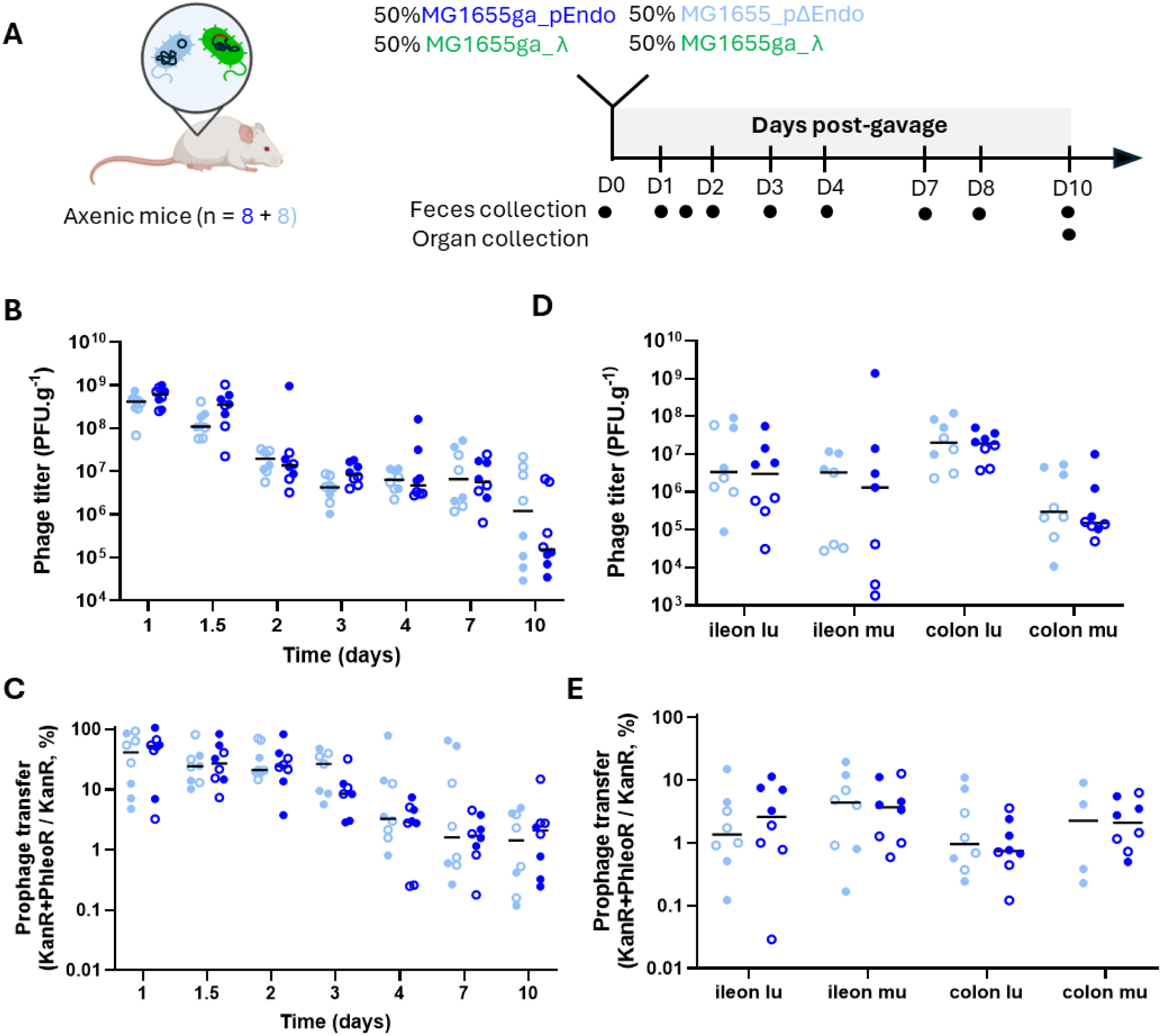
Lambda prophage induction and transfer is not increased by the E-type endonuclease. **A.** Experimental setting to test prophage-inducing ability of the E-type endonuclease producing strain. Axenic mice were orally gavaged by strain MG1655ga_λ and either MG1655ga_pEndo (n=8) or MG1655ga_pΔEndo (n=8) (2 x 10^7^ CFU of each strain). **B-C.** Phage lambda titres were determined by spot assay from feces (**B**) and from indicated gut sections (**C**; lu, luminal; mu, mucosal). **D-E.** Acquisition of Ur_lambda::ble was assessed by spot assay on selective medium, estimating the proportion of lysogenized among susceptible bacteria, in feces (**D**) and in indicated gut sections (**E**; lu: luminal; mu: mucosal). Data with median (horizontal bars) are displayed from two independent experiments distinguished by full and empty circles.

Proteomic analysis of fecal samples collected at days 0 and 10 from mice receiving MG1655ga_λ and MG1655ga_pEndo confirmed detectable expression of E-type endonuclease colicin, although the abundance rank of the protein was lower in feces than in supernatants from overnight cultures (**S3 Table**).

Induced phage lambda titres peaked at 10⁸–10⁹ PFU.g^-1^ of feces one day post-gavage and then gradually declined in the following days, consistent with previous observations using isogenic strains [33] (**Fig 6B**). Proportion of new lysogens exceeded 50% at day 1, before to decrease over time (**Fig 6C**). No significant differences in phage titres or prophage transfer were detected in feces or gut sections between MG1655ga_pEndo and MG1655ga_pΔEndo (**Figs 6B-E**). Nevertheless, we noticed that phage levels were slightly higher with MG1655ga_pEndo compared to MG1655ga_pΔEndo, reaching 2.5-fold difference in average at day 1.5 albeit non-significant (p=0.0830, Mann-Whitney test) (**Fig 6B**). This observation suggests that interactions between the colicin-producing strain and the lysogen could be transient and possibly more frequent in the first days following colonization. However, an independent experiment performed during 48 h did not confirm this trend (**S6A-B Figs**), leading us to conclude that colicin-mediated prophage induction did not occur at a scale detectable in these experimental settings.

The bacterial loads of the two strains in competition revealed a contrasted situation compared to the *in vitro* experiments. Lambda lysogens were maintained at high titres over 10 days while *in vitro* they were outcompeted in 6 h (**S7B Fig**). Nevertheless, at the end point, we observed a discrepancy between the two replicates. In one experiment a significant reduction was observed for the competition with the strain MG1655ga_pEndo compared to strain MG1655ga_pΔEndo (full dots, **S7A-B Figs**) but not in the other (clear dots, **S7A-B Figs**). The same discrepancy was observed for the bacterial loads recovered from intestinal organs (**S7C-D Figs**). This heterogeneity could be due to subtle differences in the initial ratio of the two strains or in the early stages of the competition.

## Discussion

Intestinal bacteria are often polylysogens but factors triggering prophage induction remain elusive. Likewise, consequences of prophage induction on bacteria-bacteria competition are poorly understood. Here we assessed the role of bacteriocins on prophage induction and bacterial competition.

In a collection of 1,768 *E. coli* fecal isolates from a cohort of 647 children, we found that 300 strains produced in pure cultures a soluble toxic compound. Investigating further a random set of 30 of these strains, we identified a total of 74 bacteriocin-encoding genes in 28 of them, which represents a ratio of 2.6 bacteriocin gene per strain. This is in agreement with previous studies estimating the presence of bacteriocin-encoding genes in human microbiomes [15]. Among the 74 genes, those encoding for colicin Ia and E1, and microcins V, M and H47 were the most prevalent, as expected [3,15]. However, we noted that 20% (6/30) of the strains carried an endonuclease colicin gene, a frequency that is 6-fold higher than previously estimated [15]. This could be due to their polymorphism and the consequent failure to detect all of them by PCR screening compared to our sequencing approach.

We selected an isolate that encodes two bacteriocins, an E-type endonuclease and the microcin Mcc1229, each carried by a dedicated plasmid, to decipher their relative impact on prophage induction *in vitro*. Both were independently capable of inducing the *E. coli* model prophage lambda and to promote the formation of new lysogens. More surprising was their ability to induce a broad range of prophages encompassing several virus genera, showing that bacteriocins have potentially a broad impact on lysogens residing in the gut microbiota. These results also demonstrate the benefit to combine plaque assays, growth monitoring and virome sequencing for investigating prophage induction, despite each of these approaches having its own constraints. Plaque assays offer a direct measure of released infectious phage particles, despite being limited by the choice of the indicator strain. Growth monitoring proved occasionally informative but was overall a poor predictor of prophage induction. Finally, virome sequencing of culture supernatants is time-consuming but provide a more comprehensive identification of the range of induced prophages but is dependent on total phage amounts produced.

The microcin Mcc1229 was initially identified from supernatants of human-associated *E. coli* isolates screened for their ability to enhance Shiga toxin production [14,34]. Although its precise mode of action is unknown, Mcc1229 activates RecA [14], like all endonuclease colicins. Therefore, in this study we identified only bacteriocins associated with the canonical pathway of prophage induction. However, we noticed that the supernatants from isolates 1C1 or 19G3 that putatively contain MccB17, which also activates RecA [14], did not induce the *Bievrevirus* bv_4C10 prophage. This highlights a limitation of our approach linked to the choice of a single lysogen for the first line of screening, that may be resistant to some bacteriocins. Beyond, we cannot exclude that additional bacteriocins such as MccB17 may trigger bv_4C10 excision but kill the host before virions are formed. Then, the outcome of the screen is balanced by the relative kinetics of prophage induction and toxin cytotoxicity, which depend on prophage type, bacterial fitness and susceptibility to the bacteriocin. Alternatively, some bacteriocins may target molecular machineries that are essential for phage replication and virion assembly.

*In vitro* competition assays between a bacteriocin-producing strain and a lysogen, performed in liquid and solid conditions, showed that both bacteriocins drive prophage induction that leads to a reduction of the lysogen density. Interestingly, boost on prophage induction were more pronounced in the solid condition for the microcin (**S5A-B Figs**). We speculate that the endonuclease diffuses and acts even when diluted in a well-mixed medium whereas the microcin may require a high local concentration to manifest its activity. We noticed in particular that MG1655ga_pEndo outcompeted MG1655_λ but not its non-lysogen counterpart (MG1655ga) in solid condition, whereas in liquid condition both strains were equally affected (**Figs 5F-G**). By contrast, the weak susceptibility of a lambda lysogen towards the E-endonuclease colicin in mice gut is congruent with the abundance of lysogens in the intestinal microbiota.

One of the main consequences of these competition assays was the massive generation of new lysogens among the E-type endonuclease colicin producers. However, *in vivo,* we noted that the proportion of new lysogens progressively decreases, suggesting that strains combining the prophage along with the pEndo or the pΔEndo plasmid are counterselected in the gut (**Fig 5D**).

Competition assays in the gut of mice confirmed that this environment is favourable to the induction of phage lambda [33] and that presence of a strain expressing the E-type endonuclease colicin may slightly but non-significantly increase this induction at day 1 following gavage (**Fig 6**). We hypothesize that the lack of continuous increase of prophage induction by the E-type endonuclease could be due to a weak local concentration of bacteriocin and/or a spatial distance between the two populations, overall reducing the probability of encounter. Alternatively, localized induction events in the gut may fall below the threshold of detection.

Previous studies found no measurable benefit for bacteriocin-producing strains over susceptible competitors [35], while some reported an increased fitness, especially in inflammatory contexts [36–38], as shown for example with microcin M and H47 produced by the gut isolate *E. coli* Nissle [37]. Therefore, it is possible than an inflamed gut represents a more appropriate context to detect the role played by bacteriocins in the competition between bacteria.

Overall, multiple strains carrying bacteriocins and/or prophages cohabit in a dynamic equilibrium in the gut. We propose that perturbations, whether local or systemic, increase bacteriocin levels, thereby enhancing prophage induction. However, the rise in prophage activity may have a limited impact on microbiota structure, as lysogeny buffers prophage propagation.

## Methods

### Bacterial strains and media

Human *E. coli* isolates originate from fecal samples of one-year-old children included in the COpenhagen Prospective Studies on Asthma in Childhood 2010 (COPSAC_2010_) [28]. In this study, all strains are named based on their position in 96-well plates kept at -80°C. All additional strains used for this study are listed in **Table 2**. The two *E. coli* strains used to perform spot-on-lawn assays and phage titrations are MG1655 *hsdR* (MAC1403) and strain C. None of them carry bacteriocin genes.

Unless indicated otherwise, strains were cultured in LB Lennox (5 g.L^-1^ NaCl) at 37 °C with shaking (180 rpm). The following antibiotics were used as necessary: chloramphenicol (30 µg.mL^-1^), phleomycin (10 µg.mL^-1^), kanamycin (50 µg.mL^-1^).

### Spot-on-lawn assays

5 mL of Top agar (10 g bacto-tryptone, 5 g NaCl, 4.5 g agar for 1L) pre-heated at 55°C mixed with 100 µL of overnight (O/N) cultures of strains MG1655 *hsdR* or C strains was poured onto LB Lennox agar plates. Then, 5 µL of O/N cultures of the test strains were spotted onto the lawn. Plates were incubated at 37°C O/N before observing ZOI.

### Identification of strains spontaneously releasing virions

To identify strains spontaneously releasing phage particles, 15 µL of filtrated supernatant (0.2 µm porosity, 13 mm diameter) of O/N cultures were streaked on LB agar plates and then covered by Top agar supplemented with 5 mM CaCl_2_, 10 mM MgSO_4_ and maltose 0.2%. MG1655 *hsdR* or C strains were used as indicator strains.

### Plasmid extraction, library preparation and sequencing

Plasmids were isolated using QIAprep Spin Miniprep kit (#27106). Extracted DNA was purified using AMPure XP beads (Beckman #A63882), recovered in 120 µL H2O and sonicated using Covaris (DNA 300bp) in Covaris microTUBE (Covaris, #520045). Preparation of the samples for paired-end sequencing on an Illumina NextSeq500 (2x50 bp) was conducted using Invitrogen TM Collibri TM PS DNA Library Prep Kit (Invitrogen, #A39003024) for Illumina and following manufacturer instructions.

### Plasmid assemblies and bacteriocin gene investigations

Read quality trimming was performed using Trimmomatic (options ILLUMINACLIP:TruSeq3-PE.fa:2:30:10 LEADING:3 TRAILING:3 SLIDINGWINDOW:4:20 MINLEN:30). Paired and unpaired trimmed reads were assembled with metaSPAdes from SPAdes v3.15.5. Colicins and microcins automatically annotated by Prokka v1.14.5 were recovered [40].

To complement the automatic detection, sequences of all known colicins and microcins already characterised in *Enterobacteriaceae* were collected (see **S1 Table** for UniProt or NCBI accession numbers) and used to build an updated in-house database (n=41, as of 11/2024, available online, see data accession) [16,41–43]. For colicins E4, E5 and E8, verified UniProt sequences (accessions Q47109, P18000 and P09882, respectively) were used; these correspond to partial proteins including the C-terminal catalytic domain (**S1 Table**).

All translated sequences encoded in the plasmid contig dataset were then aligned to this database using blastx. Due to the short size of microcin sequences, blastx parameters were adapted to allow screening of low-size proteins (<60 aa) with the following options: Word size:2; matrix: PAM30; Disabled SEG filtering algorithm; E-value: 0.01. Results were processed, filtering out hits having below 80% of identity and 90% of protein coverage. Annotated GenBank files were manually enriched with all the identified bacteriocins, including MccM-encoding genes missed by the Prodigal gene caller (files available online, see data accession). Positions of the bacteriocin genes on each contig are enlisted in **S2 Table**.

### Detection of microcins H47 and M by PCR

Previously reported primers and PCR settings were used to detect the presence of microcins H47 and M, using *E. coli* Nissle 1917 as a positive control [15].

### Prophage induction tests

For all assays, bacteriocins were prepared freshly from O/N liquid cultures by centrifugation (7 min – 6000 g, at 4°C) followed by filtration (0.45 μm) of the supernatants that were kept on ice until use.

O/N lysogen cultures were refreshed by 100-fold dilution in LB and incubated until OD_600nm_ = 0.2 was reached. Then, 100 μL of culture were set into 96-well plates along with 100 μL of the bacteriocin-containing supernatants. Based on the observation that addition of such a large proportion (50% of final volume) of nutrient depleted supernatants negatively impacted bacterial growth and prophage induction, supernatant of an O/N MG1655 culture was used as a negative control for spent medium. MmC was used as positive control of prophage induction (1 μg.mL^-1^). Microplates were closed with a semi-permeable film (Brand, #4TI-0516/96) and incubated 3 h in a Tecan plate reader at 37°C. OD_600nm_ was measured every 15 minutes after orbital shaking (240 sec – amplitude 5 mm).

To recover phage supernatants, after the 3 h growth and colicin treatments, microplates were centrifuged (524 x g, 5 min at 4°C) and filtrated at 0.2 µm using NucleoVac 96 Vacuum Manifold (Macherey-Nagel, #10035103) and AcroPrep™ Advance 96-well filter plates (PALL, #8019). The resulting supernatants were 10-fold serially diluted in SM buffer (100 µM NaCl, 50 mM Tris-HCl at pH 7.5 and 10 mM MgSO_4_). Five mL of top agar, supplemented with 0.2% maltose, 10 mM MgSO_4_ and 5 mM CaCl_2_ and pre-heated at 55°C were mixed with 100 µL of O/N culture of the C indicator strain and poured onto LB agar plates. Plates were allowed to dry a few minutes, and serial dilutions (down to 10^-6^) were spotted in 10 µL drops glided on the plates. Plates were then incubated at 37°C O/N and viral titres were deduced from plaque counts.

For each test, relative phage yield was determined as the ratio between the phage titre of the lysogen treated with a bacteriocin supernatant, and the mean titre of the same lysogen incubated with spent MG1655 supernatant in triplicate. Bacteriocin supernatants were considered to increase prophage induction when at least 2/3 of replicates showed titres 5-fold higher than the negative control.

### Cloning and transformation

pEndo and pMcc1229 plasmids derive from p1_1E12 and p2_1E12, two plasmids identified in isolate 1E12, respectively. A kanR cassette amplified with primers OPM60-OFL368 from plasmid pACYC177 was inserted by blunt-end ligation into p1_1E12 cut with *Pml*I. Similarly, a kanR cassette amplified using primers CH1-CH2 was cloned into p2_1E12 following digestion of the insert and the vector with *Bam*HI (**Table 3**). Ligations were transformed by heat-shock (37°C – 5 min) into RbCl-competent JM105 cells, and transformants were selected on kanamycin.

Deletions in bacteriocin cluster were then performed as follows: pΔEndo results from the digestion of pEndo by *Dra*I followed by self-ligation. In the case of pΔMcc1229, the plasmid was digested with *Bmt*I and *EcoR*I, and the resulting overhangs were blunted using DNA Polymerase I Large (Klenow) Fragment (NEB, #M0210) prior to ligation. Ligations were transformed as above and transformants were selected on kanamycin. Deletions in pEndo and pMcc1229 were verified by PCR using CH22-CH23 and CH14- CH15 primers, respectively (**Table 4**). Whole plasmid sequencing for pEndo, pMcc1229 and their respective controls confirmed the constructions and the absence of unintended mutations.

**Table 4.**
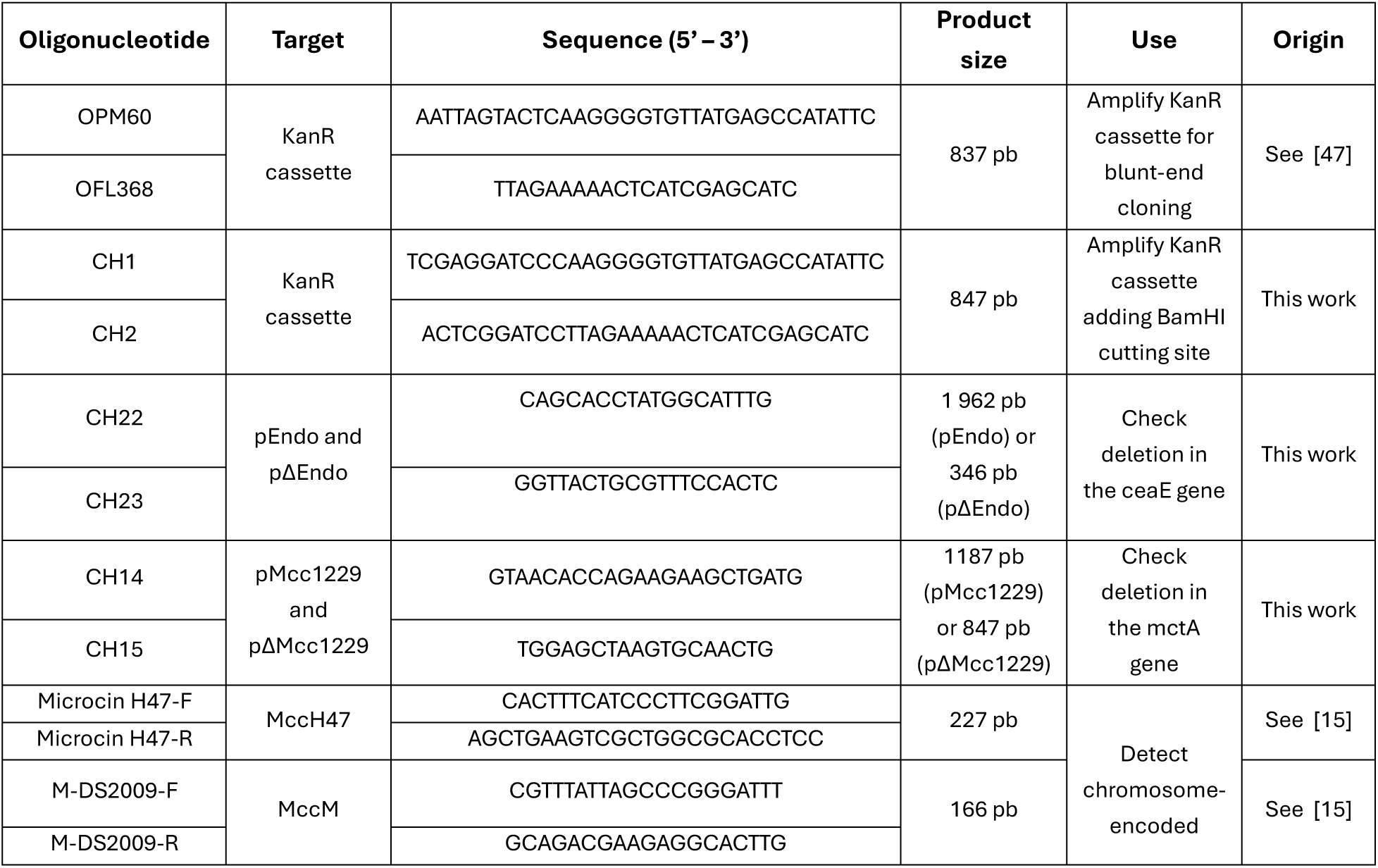
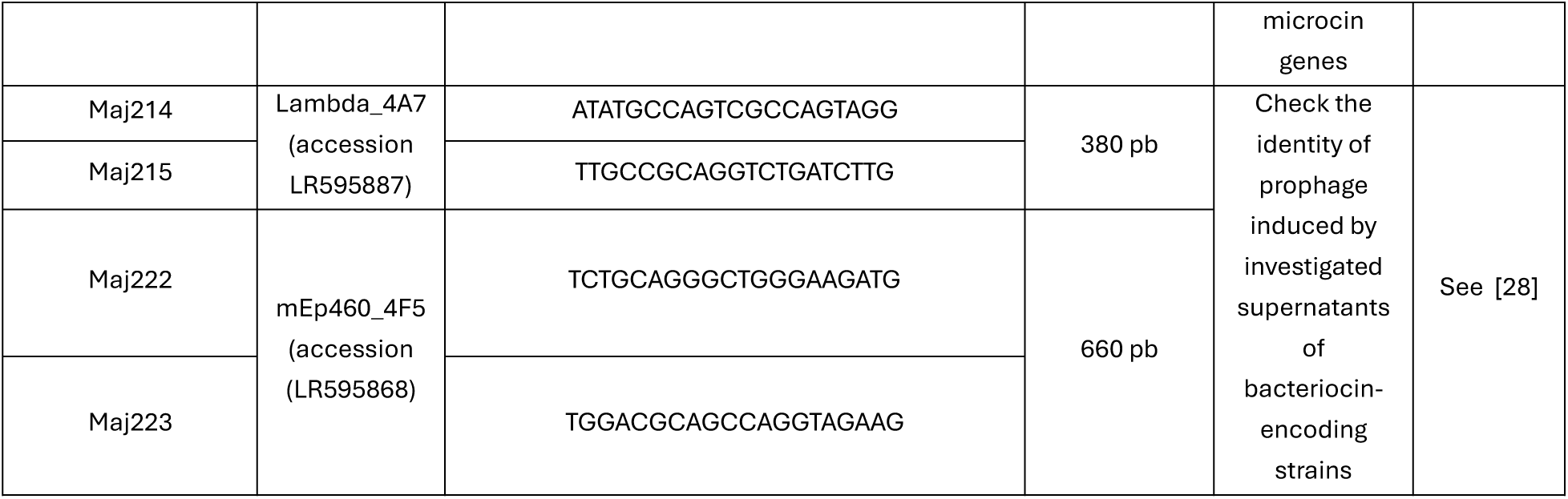
Primers used in this study.

Plasmids were then transferred into the strains to be tested, MG1655 and MG1655ga, by chemical transformation. To do so, corresponding cultures were grown to early exponential growth (OD_600nm_=0.18-0.22), pelleted and permeabilised by resuspension in 1 mL of ice-cold 100 mM CaCl_2_. Following a 4 h incubation, 1-10 µL of plasmid DNA was added to 100 µL of competent cells. Cells were incubated on ice for 20 min, subjected to heat shock (37°C – 5 min) and allowed to recover in SOC medium for 1 h. Transformants were then selected on LB agar supplemented with kanamycin. The resulting strains are listed in **Table 2** and plasmid constructions in **Table 3**.

### Virome sequencing and analyses

The same conditions used for prophage induction tests in small cultures were repeated in 100 mL. At OD around 0.2, 100 mL of O/N culture supernatants (from MG1655 or MG1655_pMcc1229 strains), filtered at 0.45 µm (Fischer) were added to each 100 mL exponentially growing culture, and incubated at 37°C 3 h. Bacteria were then pelleted (5000g, 15 min, in four 50 mL tubes per culture), supernatants were recovered and filtered. Next, NaCl was adjusted to 0.2 M and PEG8000 was added to 10% of the total supernatant volume. After PEG solubilisation and O/N incubation at 4°C, the phage pellet was recovered by centrifugation (5000g, 15 min) and resuspended in DnaseI buffer (500 µL for each culture). The phage suspensions were treated for 30 min at 37°C with 1 µL of DnaseI (Ambion), and next, phage proteins were removed with two successive phenol-chloroform-isoamylalcohol (25:24:1) extractions, followed by a chloroform-isoamylalcohol (24:1) extraction. Finally, DNA was precipitated after addition of 1 µL glycogen, 0.3 M potassium acetate and 0.6 vol isopropanol. The DNA pellet was washed with ethanol 70%, and air dried. For the 4C9 viromes, where insufficient DNA had been recovered for sequencing, the Genomiphiv3 kit was used to amplify DNA sequences. All samples were sequenced at Eurofins (Germany), with Illumina technology and 2x150 nt paired ends. Read cleaning and assembly were performed using SPAdes and default parameters.

To estimate the proportion of each phage genome in the viromes, reads were mapped back onto the assembled phage sets of each virome, including the P2 genome. Input reads were 13 million for 4E6 viromes, 1 million for 4C9 viromes and 0.5 million for the Mcc1229-treated 2H4 virome. The control MG1655-treated 2H4 virome was heavily contaminated, so that initial proportions of the P2 and the *Jouyvirus* genomes were estimated from 140 reads mapped to P2, and 6 to the *Jouyvirus* phage genome. Mapping was conducted with the bwa-mem2 software, version 2.2.1, and default parameters [44]. The resulting SAM file was converted to BAM and sorted by read name using samtools version 1.9, before filtering out the low-quality alignments using msamtools filter version 1.1.3 [45]. Alignments >80 bp with >95% identity over >80% sequence length were selected, and only the best alignment for each read were kept. The coverage and depth of each genome for each sample was determined using msamtools coverage. Relative coverage of each phage genome compared to P2 was then computed and compared for the microcin-treated (MG1655_pMcc1229 supernatant) and MG1655-spent medium treated control.

### Competition assays

O/N bacterial cultures grown with respective antibiotics were washed in LB medium twice (10 min, 6000g at 4°C). Then, bacterial cultures were refreshed by 1:100 dilution in LB and incubated at 37°C until OD_600nm_ reached 0.3. For a given experiment, MG1655ga_pEndo and MG1655ga_pΔEndo (or pMcc1229 and pΔMcc1229) were mixed with the MG1655ga_λ lysogen, in a 1:1 ratio. Pairwise cultures were then plated immediately to assess that both strains of each pair were initially distributed in equal quantities.

Bacterial competition was explored in two conditions. First, 5 mL of the mix was incubated in tubes at 37°C with shaking at 180 rpm (Liquid condition). In parallel, 5 µL of the mixes were spotted on LB agar plates and incubated after the spot dried (Solid condition). Strains were co-cultured for 6 or 24 h. Then, to count viable cells for each population, mixes of the solid condition were cut from agar plates (using a 1 mL tip sectioned at the appropriate diameter) and resuspended in 5 mL of LB. Then, all bacterial suspensions were serially diluted in LB and spotted onto LB agar plates supplemented with kanamycin or chloramphenicol. Titration of Ur_lambda was performed as described above. The level of lysogenization of the KanR, phage sensitive strain by Ur_lambda::ble in the co-cultures was assessed by plating them on both kanamycin and phleomycin. The prophage transfer efficiency was evaluated by calculating the ratio of double resistant KanR PhleoR over KanR clones.

### Animals and ethics

Axenic C57Bl/6J, 7–8-week-old and bred at Institut Pasteur were used. Mice were randomly assigned in each group with males and females separated. All mice were housed in a germ-free animal facility and maintained in isocages in accordance with Institut Pasteur guidelines and European recommendations. Food and drinking water were provided *ad libitum*. All animal experiments were approved by the committee on animal experimentation of the Institut Pasteur and by the French Ministry of Research (APAFIS n° #26874-2020081309052574 v8) and were performed according to the legal requirements.

### Colonization of axenic mice

Strains for gavage were inoculated in 5-mL LB medium with appropriate antibiotics. Cultures were washed twice in fresh LB to remove antibiotics, phages and toxins (centrifuge 6000g, 10 min at 4°C). To identify the impact of the strain producing the E-type endonuclease on prophage dynamics *in vivo*, the two bacterial strains were gavaged orally together. Mice received 200 µL of bacterial culture with 10^7 C^F^U^ of each bacteria in sucrose sodium bicarbonate solution (20% sucrose and 2.6% sodium bicarbonate, pH 8). Mice were monitored and weighted daily.

At given time points, fecal pellets were collected and transferred in 1.5-mL tubes, weighted and resuspended in 1 mL of PBS. 100 µL of these suspensions were serially diluted in LB medium and spotted on LB plates with appropriate antibiotics to estimate bacterial loads. To assess levels of phages, the remaining suspension was centrifuged (6000g, 10 min at 4°C), and 100 µL of the upper supernatant was carefully taken, serially diluted in TN buffer (10 mM Tris-HCl, 150 mM NaCl, pH 7.5) and plated on bacterial lawn of C strain as described above.

The experiments were ended at day 10 (2 experiments; each with 2 x 4 mice) or at day 2 (1 experiment; 2 x 4 mice). Following cervical dislocation, intestinal sections of ileums and colons were collected and luminal and mucosal parts were separated. The luminal part was recovered by squeezing the intestinal tube with the back of a scalpel to recover the content that was subsequently homogenized in 1 mL of PBS. The remaining tissues corresponding to the mucosal part were washed in 10 mL of PBS before being transferred into a new tube with 2 mL of fresh PBS and then homogenized using the gentleMACS^TM^ OctoDissociator (Miltenyi Biotec). Serial dilutions in LB were plated on LB plates containing appropriate antibiotics.

### Liquid chromatography mass spectrometry of proteome extracts

Fecal samples (at least 20 mg wet weight in total) collected at day 1 and day 10 (n = 2 for each timepoint, from two independent experiments) were resuspended in 1 mL PBS, homogenized and then centrifuged (3 min – 5000 g – 4°C). Pellets and supernatants were processed separately as follows including a positive control of 1 mL of an MG1655ga_pEndo O/N culture supernatant centrifuged (5000g, 6 min at 4°C°) and filtered at 0.45µm.

Proteins were digested using an adapted S-Trap™ protocol [46]. Samples were first solubilized in SDS lysis buffer (4% SDS, 50 mM Tris-HCl, pH 7.5), homogenized on a Covaris E220 sonicator using Adaptive Focused Acoustics (AFA) technology and clarified by centrifugation. Proteins were then reduced with TCEP and alkylated with iodoacetamide. Samples were acidified, diluted in S-Trap binding buffer (100 mM AMBIC in 90% methanol/10% water) and loaded onto S-Trap columns. After washing, proteins were digested on-column overnight at 37 °C with sequencing-grade trypsin. Peptides were sequentially eluted, pooled, dried by vacuum centrifugation, and stored at −20 °C until LC–MS/MS analysis.

All peptide fractions were resuspended in 0.1% formic acid immediately before LC–MS/MS injection. Peptide analysis was performed on a timsTOF Ultra mass spectrometer (Bruker) coupled to a nanoElute LC system (Bruker). Peptides were loaded onto a PepSep Twenty-five Series column (75 µm i.d., 1.5 µm particle size, Bruker). Chromatographic separation was achieved with a linear gradient at a flow rate of 250 nL.min^-1^ and a column temperature of 50 °C. CaptiveSpray source was operated with a capillary voltage of 1,600 V, a dry gas flow of 3.0 L.min^-1^, and a temperature of 200 °C. MS data were acquired in dia-PASEF (data-independent acquisition parallel accumulation–serial fragmentation) mode. The full m/z range from 100 to 1,700 Th was covered using optimized isolation windows aligned to ion mobility values spanning 0.64–1.45 Vs/cm² (1/K₀). Each dia-PASEF cycle comprised multiple MS/MS scans with a total duty cycle of ∼1.1 s. Collision energies were ramped linearly as a function of ion mobility, following the manufacturer’s recommendations.

Spectra were searched against a UniProt *E. coli* database containing the canonical and reviewed protein sequences (4,414 entries the 2026/01/28) to which the WT and truncated colicin sequences were added using Spectronaut ver. 20.3 software. Search criteria included carbamidomethylation of cysteine as a fixed modification, oxidation of methionine and acetylation (protein N terminus) as variable modifications and trypsin specificity with up to two missed cleavages was allowed. For other parameters, default settings were used. A false discovery rate (FDR) cutoff of 1% was applied at the peptide and protein levels.

### Data accession

Plasmidomes (raw read data and assembled plasmid contigs) of 24 *E. coli* bacteriocin producers, as well as the in-house colicin and microcin database, were published in Data INRAE (https://doi.org/10.57745/FVPLLS). Raw read data of the virome of the lysogens *E. coli* 2H4, 4E6 an 4C9 obtained after/without bacteriocin treatments were deposited in the European Nucleotide Archive (ENA) at EMBL-EBI under accession numbers ERR16144120/ERR16144121, ERR16144122/ERR16144123 and ERR16144124/ERR16144125, respectively (study: PRJEB106466). Bacteriocin-inducible *Escherichia* phages Josas_2H4, jv_4E6, ue_4E6 and ue_4C9 assembled and annotated genomes were deposited in the ENA at EMBL-EBI under accession numbers OZ418229, OZ418230, OZ418234, and OZ418233, respectively (study: PRJEB106466).

The mass spectrometry proteomics data have been deposited to the ProteomeXchange Consortium via the PRIDE partner repository with the dataset identifier (in process).

## Acknowledgements

We thank BBH and DynPhage team members for discussions and especially Elisabeth Moncaut, Benoît Lerouge, Camille Sivelle, Céline Mulet and Julien Lossouarn for their help in some experiments. We acknowledge M. Matondo and E. Bouscasse from the MSBio Proteomics Core Facility at Institut Pasteur (equipment partially funded by DIM1Health 2023 programme SLIM) as well as members of the Centre for Gnotobiology of Institut Pasteur. We also thank Karen Krogfeld for hosting the COPSAC strain collection and Aurélie Mathieu for the initial ZOI screening.

A CC-BY public copyright license has been applied by the authors to the present document and will be applied to all subsequent versions up to the Author Accepted Manuscript arising from this submission, in accordance with the grant’s open access conditions.

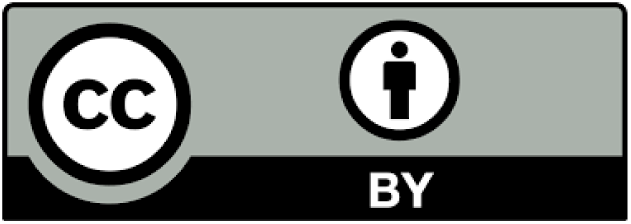

## Declaration of AI tools

Authors declared that they have used AI tools (Mistral) for language enhancement or proofreading followed by manual curation to ensure that scientific content remained intact.

## Supporting Information

### Supporting text

#### Bacteriocin content in *E. coli* isolates

The most frequent bacteriocins in our isolates were MccH47 (n=10), colicin E1 (n=8), colicin Ia (n=8), MccV (n=7), MccM (n=7) and various endonuclease colicins (n=6). Pore-forming colicins were particularly frequent since two-third of the isolates had at least one corresponding gene. Regarding the endonuclease colicins, additionally to the 6 identified clusters, strain 14C8, which has such a complete cluster, also possess a second partial sequence annotated as catalytic domain of endonuclease, which was not considered further as it could not be fully assembled.

Endonuclease colicins display a modular structure with a N-terminus T domain involved in transport, a central R domain involved in receptor binding and a C-terminus D domain containing the DNAse domain with H-N-H motif. The endonuclease colicins that we identified can be separated into two homogeneous sub-groups: a group composed of 1E12, 3A5 and 6H3 (>97.6% of amino-acid identity by pairwise comparisons) and a second group composed of 4E8, 12E9 and 14C8 (>98.6% amino-acid identity by pairwise comparison and 91.0-98.26% identity with reference colicin 1E7) (**S1B Fig**). Translocator domains (T) of these 6 colicins were all most similar to the corresponding domain of the E7 colicin, while receptor-binding domains (R) and DNAse domains (D) exhibited a high polymorphism, being more similar to colicin E2, E7 or E9 domains.

The 6 complete sequences were not perfectly identical to referenced endonuclease colicins E2-7-9, and mutations were not evenly spread. Instead, we observed modular structures of their translocation (T), receptor-binding (R) and DNase (D) domains (**S1B Fig**). This organisation is typical of polymorphic toxins, which display a huge variation with distinct domains generated through recombination [48]. This is why we chose to refer to these genes broadly as “endonuclease colicin”. Beyond, we suggest that previous analyses were established based on protein comparisons that were not representative of the diversity of these toxins, actually hiding their high variation patterns.

## Supporting figures

**Figure S1.**
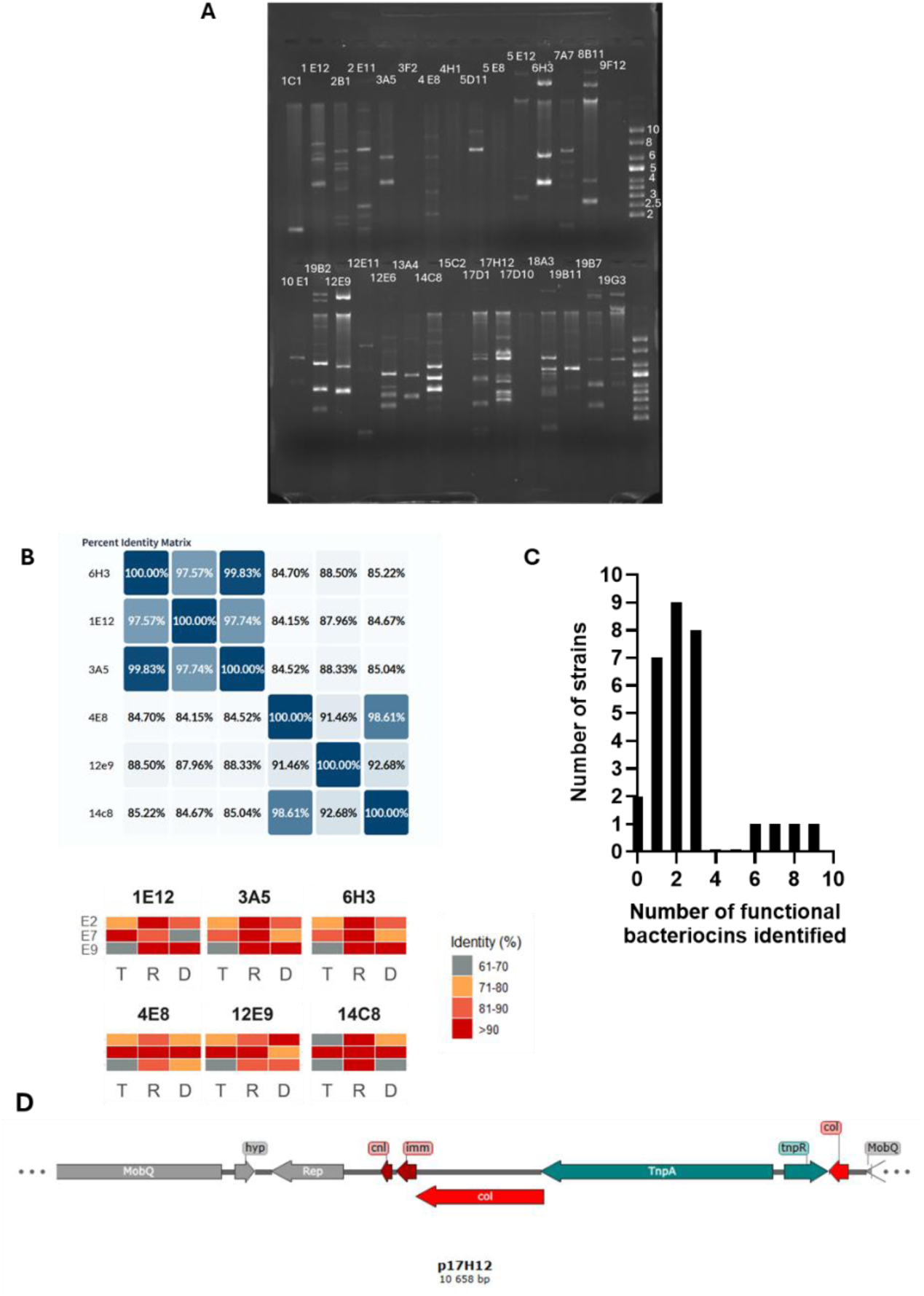
Bacteriocin screening in the 30 COPSAC isolates. **A.** Electrophoresis of plasmid content for the selected 30 COPSAC isolates (see **Fig 2A**). **B.** Number of bacteriocin genes identified per *E. coli* isolate. **C.** Domain comparison for the different E-type endonuclease colicins. Top: Identity matrix for the 6 E-type colicins (UniProt). Bottom: For each endonuclease predicted domains (Interpro 105.0) of translocation (T), receptor-binding (R) and DNase (D) (See **Annex 1**), a heatmap shows the identity (%) to the corresponding domain in endonucleases E2 (UniProt accession P04419), E7 (UniProt accession Q47112) and E9 (UniProt accession P09883). Whereas the R domain is conserved among all endonucleases, domains T and D vary: 1E12, 3A5 and 3H3 endonucleases have an E7-E9 chimeric composition, 12E9 has an E7-E2 chimera, and 4E8 is fully homologous to E7. **D.** Genetic map showing the transposon insertion within 17H12 endonuclease gene.

**Figure S2.**
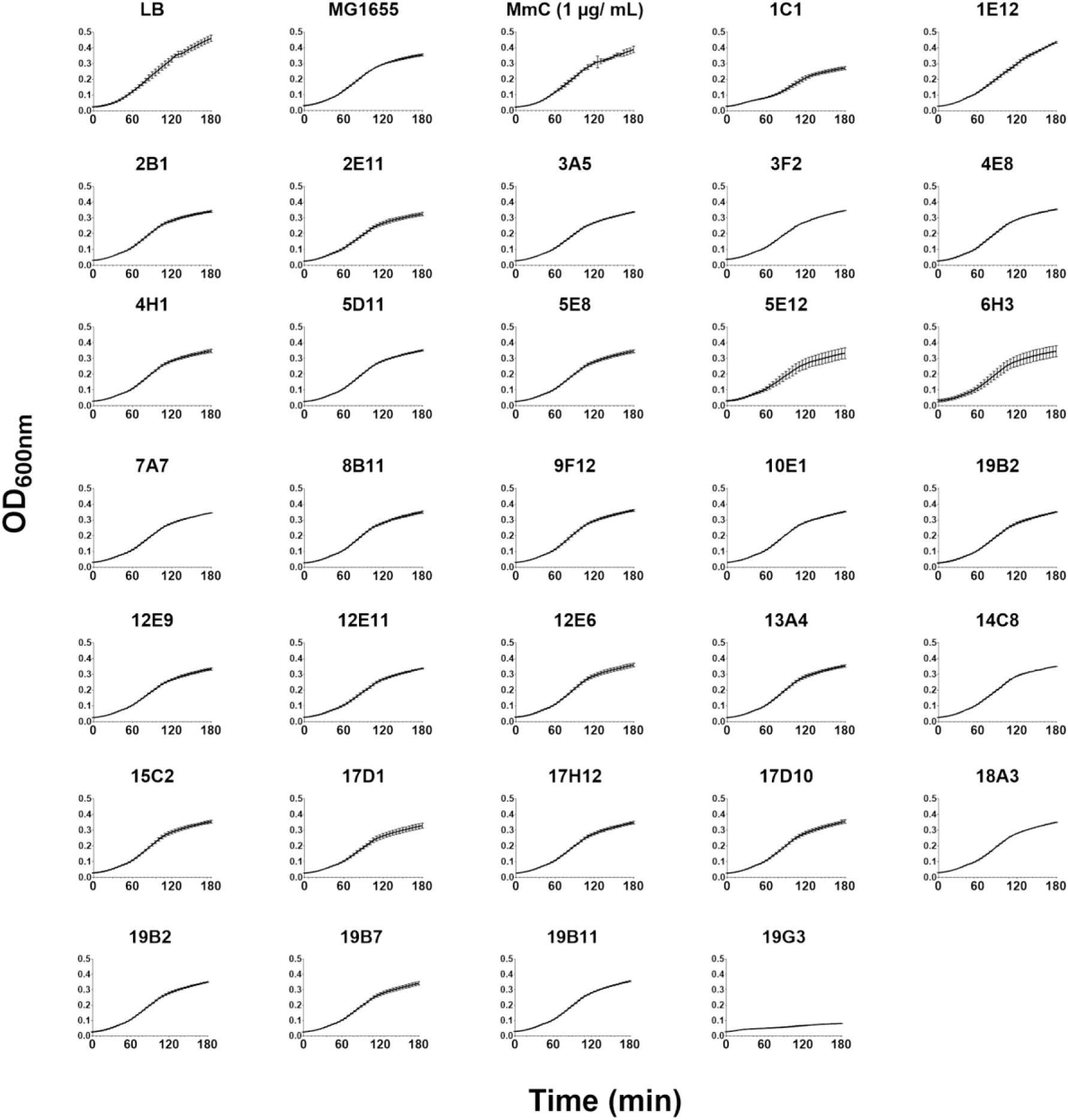
Growth curves of the *E. coli* strain 4C10 exposed to the 30 supernatants of the bacteriocin-producing *E. coli* isolates. Growth was monitored following OD_600nm_ in microplaques at 37°C using a Tecan spectrophotometer. MG1655 supernatant was used as negative control for nutrient depletion. Curves shown for each condition are average and standard deviations of 3 replicates.

**Figure S3.**
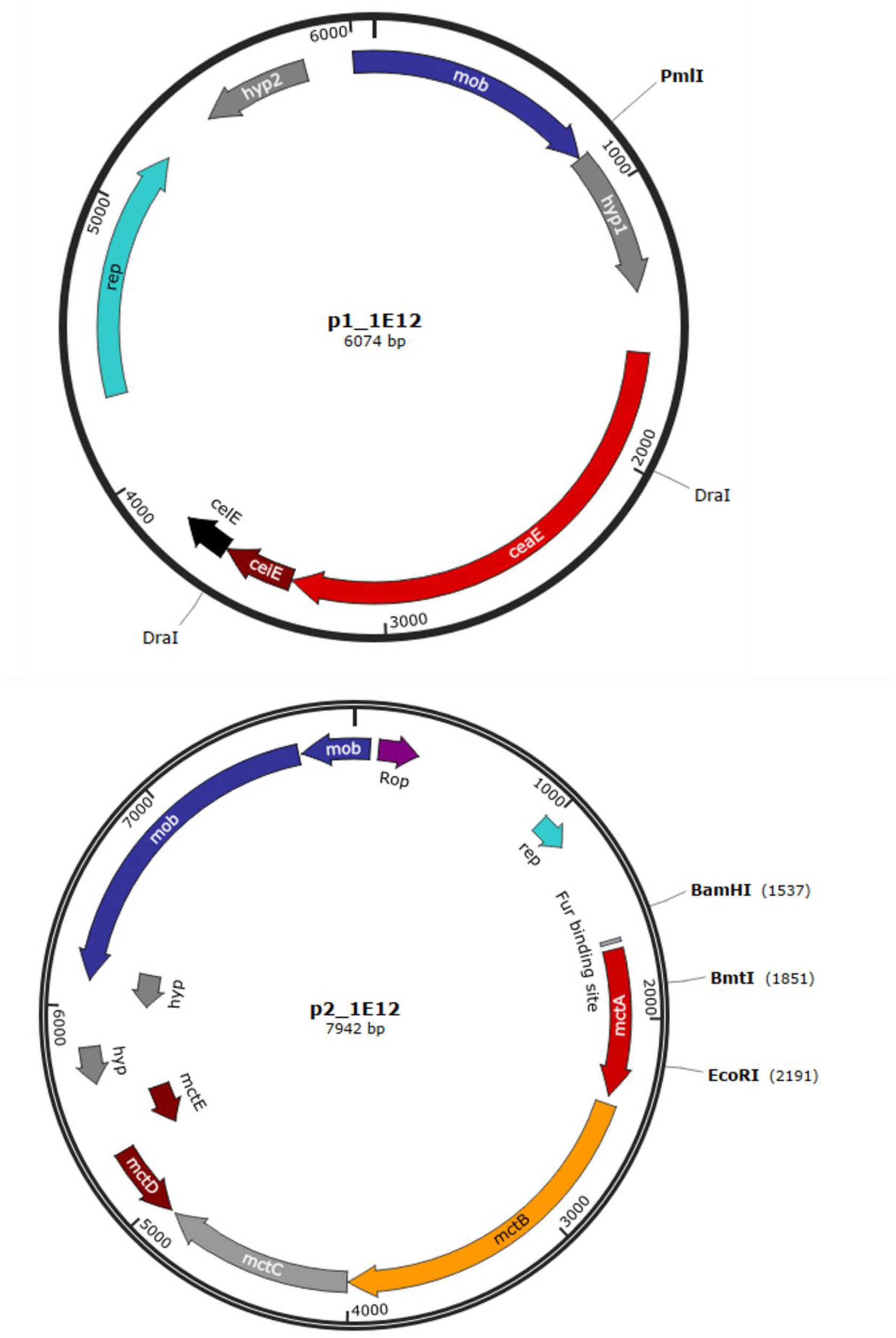
Plasmid map of p1_1E12 and p2_1E12 carrying respectively the E-type endonuclease colicin and the Mcc1229 cluster. The plasmid maps were generated using SnapGene software (from GSL Biotech).

**Figure S4.**
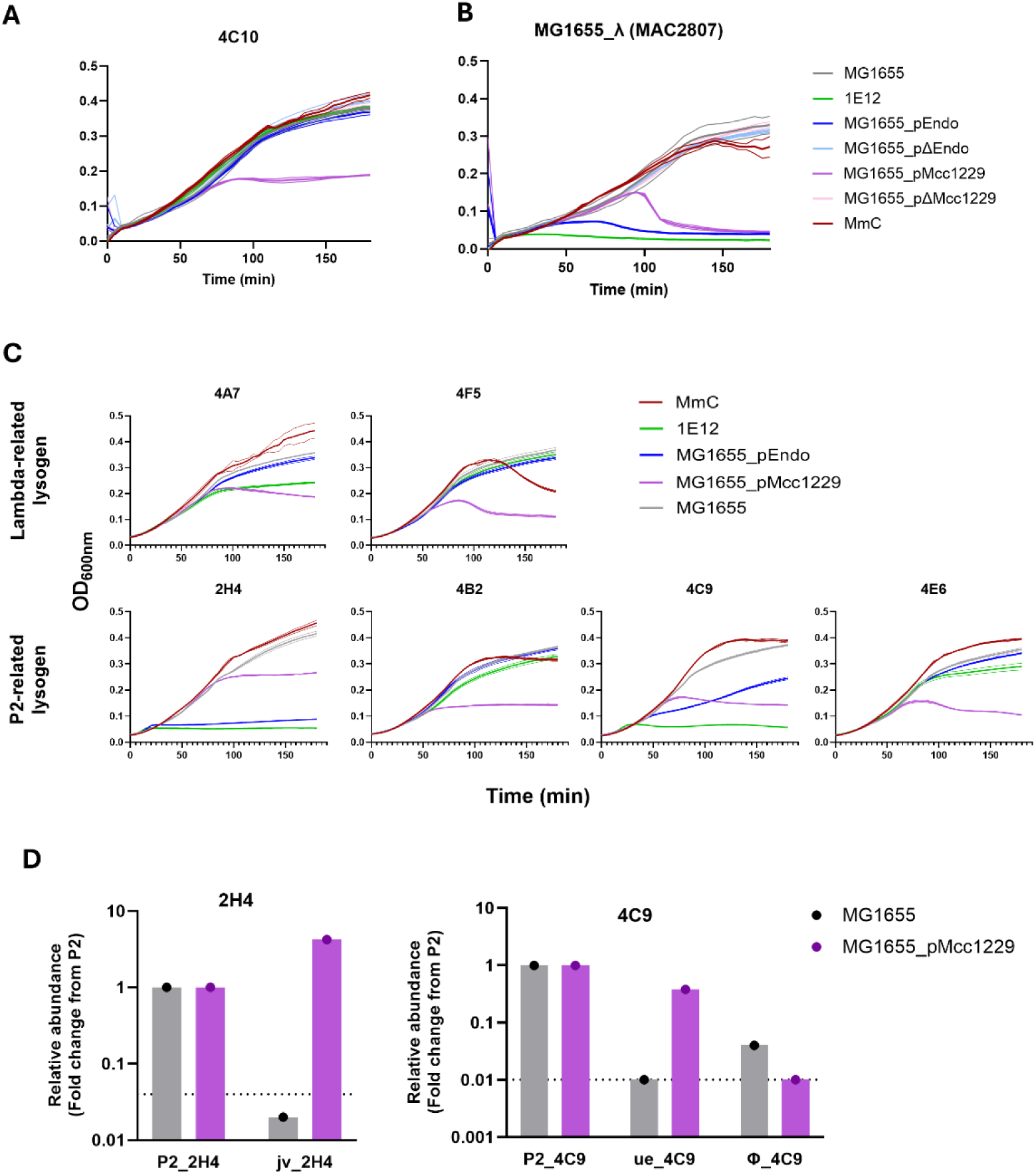
Supernatants of MG1655 carrying p_Endo and p_Mcc1229 trigger growth inhibition in different lysogens of the COPSAC collection, suggesting prophage induction. A-B. Growth curves of 4C10 (**A**) or MG1655_λ (**B**) exposed to diverse non-diluted supernatant of overnight cultures during 3 h. **C.** Growth curves for six *E. coli* lysogens from the COPSAC collection exposed to non-diluted supernatant of overnight cultures of either 1E12, MG1655_pEndo and MG1655_pMcc1229 during 3 h. These lysogens were reported to undergo spontaneous induction of lambda-related (n=2) or P2-related (n=4) prophages [28]. **D.** Virome sequencing of strains 2H4 and 4C9, treated with Mcc1229 microcin or MG1655 supernatants, revealed polylysogeny: strain 2H4 supernatant contained phage Josas_2H4 which was induced by a factor of 100 upon microcin treatment, and strain 4C9 contained phage ue_4C9 (50-fold induction) and a lambda-like phage named Φ_4C9 (uninduced). The dashed line represents the relative abundance of bacterial contigs for the lysogen treated with MG1655 supernatant.

**Figure S5.**
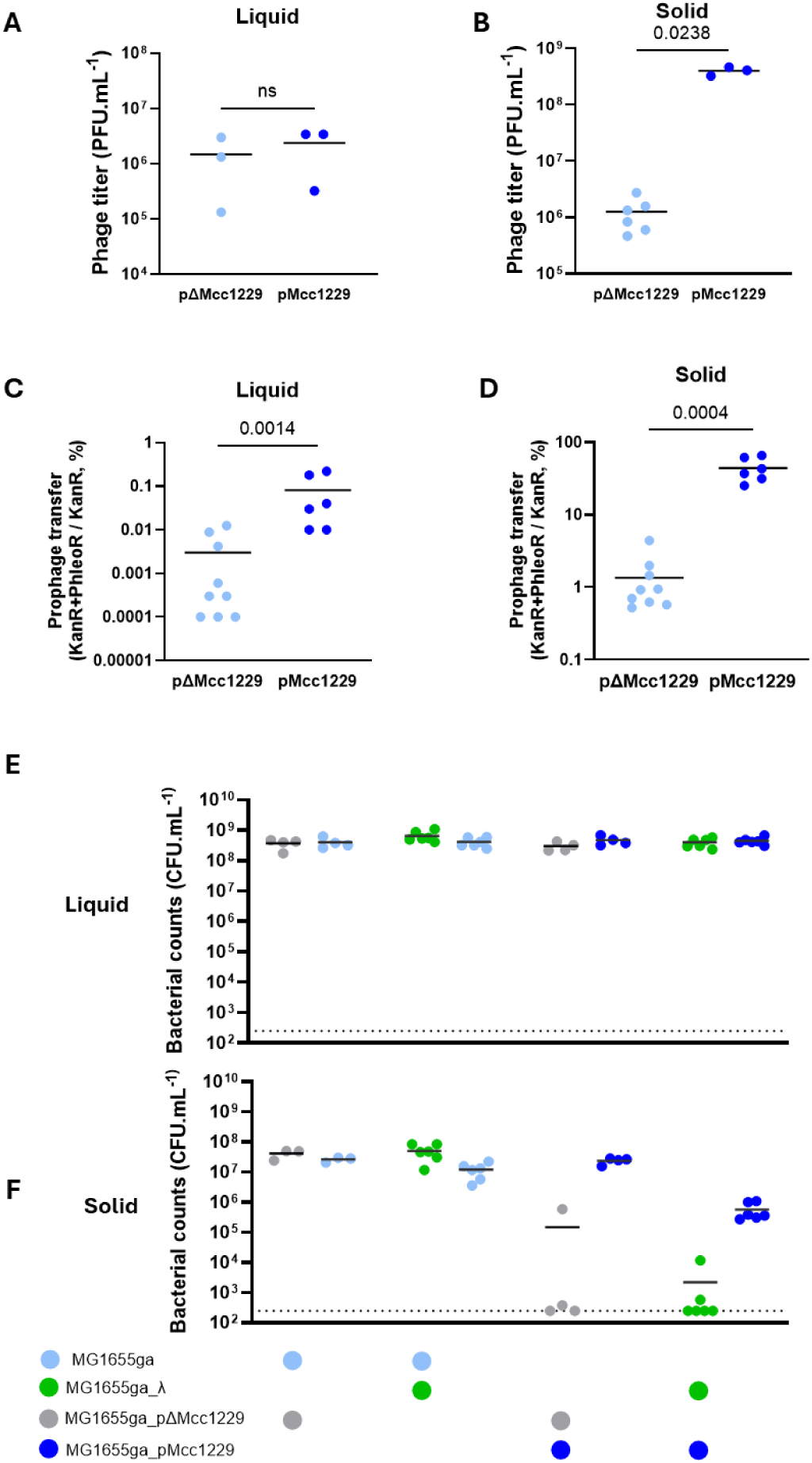
Mcc1229 induces lambda and outcompetes the lysogen in solid but not in liquid conditions, after 6 hours of co-culture. A-F. Equal volumes of exponentially growing MG1655ga_λ (CmR + PhleoR) or MG1655ga (CmR), placed in competition with MG1655ga_pMcc1229 (KanR) or MG1655ga_pΔMcc1229 (KanR), were mixed and grown in 5-ml LB broth (liquid) or on plate (solid) for 6 h. **A,B.** Free phages were titrated by spot assay on strain C. **C,D.** Acquisition of Ur_lambda::ble was assessed by spot assay on selective medium. Prophage transfer was estimated by the proportion of newly lysogenized bacteria in the MG1665ga_pMcc1229 (or pΔMcc1229) population. **E,F.** Bacterial loads were measured after 6 h of pairwise competition in liquid (**E**) or solid (**F**) conditions. Data are represented with the mean. Statistical analysis was performed using Mann-Whitney for unpaired samples.

**Figure S6.**
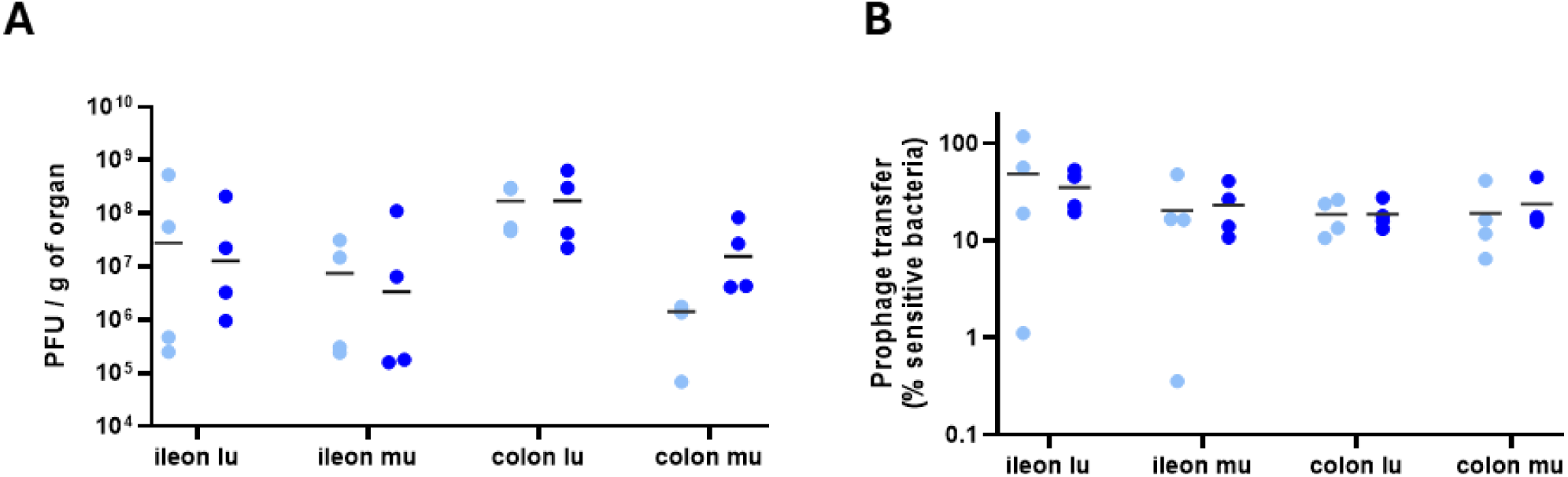
Phage titers and proportion of lysogens in gut sections at day 2. Axenic mice received MG1655ga_λ and either MG1655ga_pEndo (n=4 mice) or MG1655ga_pΔEndo (n=4 mice) by oral gavages (2.10^7^ CFU of each bacterial strain). Phage titers and new lysogens were plated from indicated gut sections (lu, luminal; mu, mucosal). **A.** Free phages were titrated by spot assay on strain C. **B.** Acquisition of Ur_lambda::ble was assessed by spot assay on selective medium as previously described. Data are represented with the median.

**Figure S7.**
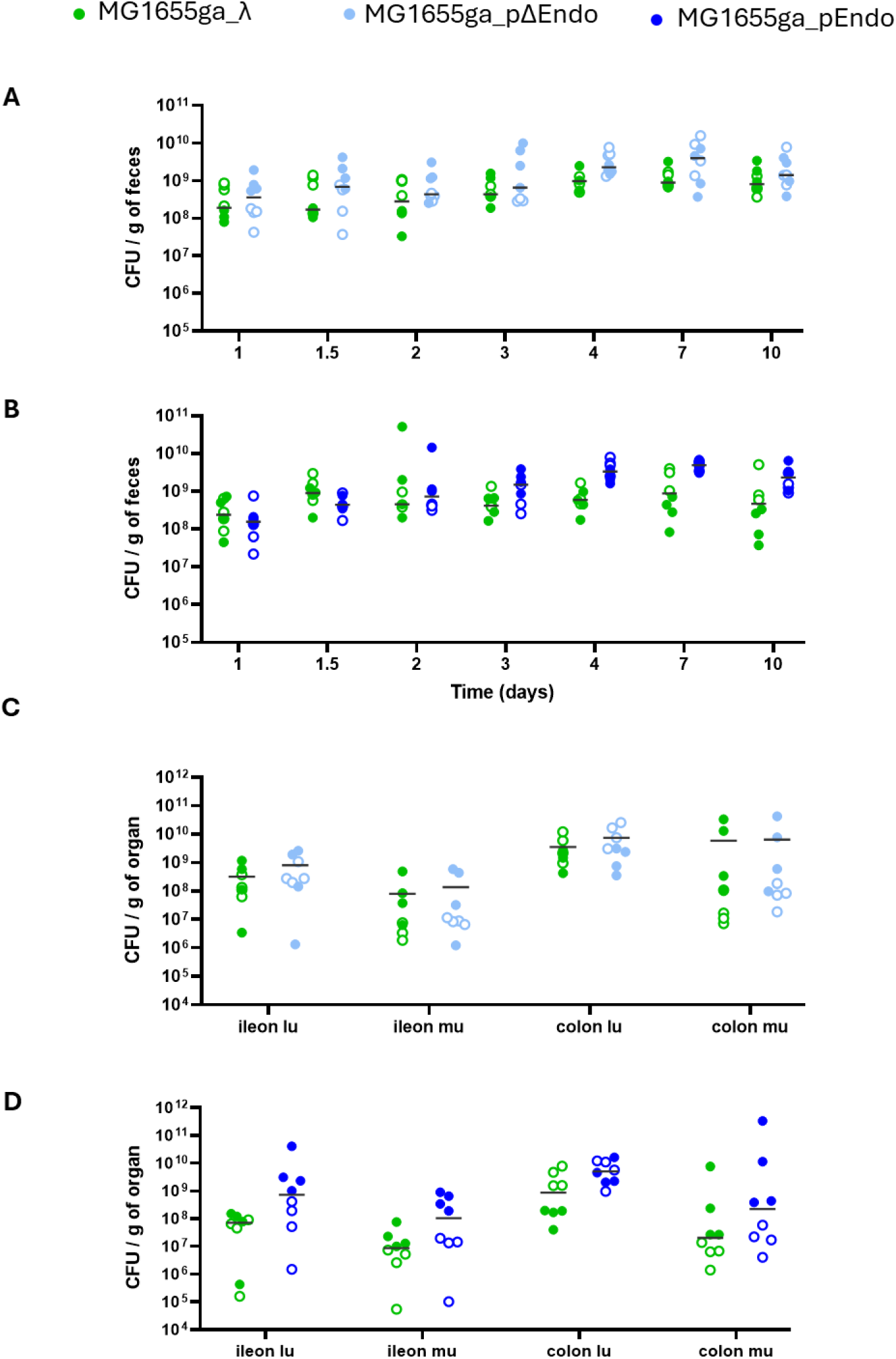
Bacterial loads in feces and gut sections from mice co-colonized with a lysogen *E. coli* MG1655ga_λ and an isogenic strain harbouring pΔEndo or pEndo. Axenic mice received MG1655ga_λ and MG1655ga_pΔEndo (**A** and **C**; n=8 mice) or MG1655ga_pEndo (**B** and **D**; n=8 mice) by oral gavages (2.10^7^ CFU of each bacterial strain). Bacterial loads were plated on selective media from fecal samples (**A,B**) and gut sections at day 10 post-gavage (**C,D**). The two independent experiments are merged, with exp.1 dataset represented by full circles, and exp.2 dataset by empty circles. Data are represented with the median.

